# PharaohFUN: PHylogenomic Analysis foR plAnt prOtein History and FUNction elucidation

**DOI:** 10.1101/2023.08.01.551440

**Authors:** Marcos Ramos-González, Víctor Ramos-González, Emma Serrano-Pérez, Christina Arvanitidou, Jorge Hernández-García, Mercedes García-González, Francisco José Romero-Campero

## Abstract

Since DNA sequencing has become commonplace, the development of efficient methods and tools to explore gene sequences has become indispensable. In particular, despite photosynthetic eukaryotes constituting the largest percentage of terrestrial biomass, computational functional characterization of gene sequences in these organisms still predominantly relies on comparisons with *Arabidopsis thaliana* and other angiosperms. This paper introduces PharaohFUN, a web application designed for the evolutionary and functional analysis of protein sequences in photosynthetic eukaryotes, leveraging orthology relationships between them.

PharaohFUN incorporates a homogeneous representative sampling of key species in this group, bridging clades that have traditionally been studied separately, thus establishing a comprehensive evolutionary framework to draw conclusions about sequence evolution and function. For this purpose, it incorporates modules for exploring gene tree evolutionary history, expansion and contraction events, ancestral states, domain identification, multiple sequence alignments, and diverse functional annotation. It also incorporates different search modes to facilitate its use and increase its reach within the community. Tests were performed on the whole transcription factor toolbox of *Arabidopsis thaliana* and on CCA1 protein to assess its utility for both large-scale and fine-grained phylogenetic studies. These exemplify how PharaohFUN accurately traces the corresponding evolutionary histories of these proteins by unifying results for land plants, streptophyte and chlorophyte microalgae. Thus, PharaohFUN democratizes access to these kind of analyses in photosynthetic organisms for every user, independently of their prior training in bioinformatics.

## Introduction

With the falling cost of next-gen sequencing technologies, the availability of genomes of new organisms is growing exponentially. This allows for the inclusion of representatives in clades that enable the reconstruction of evolutionary histories of lineages on a large scale (Rokas and Carroll 2005). However, the functional characterization of new gene sequences is not progressing at the same rate which is producing a bottleneck in the application of these organisms as models of their corresponding clades (Cantalapiedra et al. 2021).

One clade where this phenomenon has a marked impact is Viridiplantae. In 2020, the INSDC (International Nucleotide Sequence Database Collaboration) (Arita et al. 2021) contained genomes for more than 800 species of this group, but their distribution remains heavily unbalanced: while 543 correspond to angiosperms, other taxa such as gymnosperms, bryophytes or most algae are poorly represented (Kress et al. 2022). The imbalance in the functional annotation of these genomes is even more pronounced. Although organisms such as *Arabidopsis thaliana* and *Oryza sativa* have highly refined annotations, other taxa do not receive as much attention (Van Bel et al. 2018). Annotation based on these species establishes an important bias, especially since most new species do not receive computational or experimental reviews or manual curation (Van Bel et al. 2018; Kress et al. 2022). In addition, most functional annotation databases neglect entire clades of this group, as is the case for streptophyte algae, key models for understanding plant colonization of land (Huerta-Cepas et al. 2019; Leebens-Mack et al. 2019; Hanschen and Starkenburg 2020). The unequal availability of resources among species translates into a clear predominance of *A. thaliana* and other angiosperms in terms of the number of published papers (Marks et al. 2023).

In recent years, it has been demonstrated how working in an evolutionary framework facilitates the resolution of classic problems in genomics. One such example is the assignment of relationships between genes. In this context, software based on phylogenetic methods, such as SHOOT, has been found to be more accurate than those based on pairwise alignments, such as BLAST (Emms and Kelly 2022). Another instance is the annotation of genomic regions. Here, TOGA incorporates information from orthologous genes to delineate coding and non-coding regions, outperforming previous methods in terms of annotation completeness (Kirilenko et al. 2024). Finally, the use of ancestral reconstructed sequences to train Protein Language Models has been shown to improve their accuracy. In particular, this has led to the development of more meaningful epistasis representations than those produced by larger models trained without incorporating evolutionary data (Matthews et al. 2023).

In this work, we introduce PharaohFUN, a web-based tool aiming to facilitate the exploration of the evolutionary landscape of photosynthetic eukaryotes. PharaohFUN also seeks to facilitate inter-species transfer of functional information based on orthology relationships between genes. To achieve this, PharaohFUN incorporates genomic information from 167 species spanning Prasinodermophyta, Chlorophyta and Streptophyta plus 4 rhodophytes as outgroups. Thus, PharaohFUN currently supports one of the largest numbers of plant species among the software dedicated to orthology and functional transfer studies, especially for neglected non-angiosperm plant species. The phylogenetic placement of these major clades establishes a practical framework for drawing hypotheses about the evolutionary relationships between genes.

PharaohFUN integrates phylogenetic and evolutionary contexts of each gene, along with sequence and domain studies and leverages species with refined annotation. In turn, this allows users to build reliable hypotheses about the evolutionary history and function of a gene of interest in a few minutes. Unlike most existing phylogenetic software, PharaohFUN not only accelerates studies of individual genes. Through its Batch Mode, it allows simultaneous analysis of long lists of genes and sequences, facilitating large-scale evolutionary analyses. At the same time, it does not rely solely on its database species but allows the inclusion of sequences from external organisms. In addition to traditional tools for domain, sequence and protein function exploration, PharaohFUN incorporates features absent in other phylogenetic software. These include ancestral state reconstruction of gene families, protein-protein interactions and literature annotation.

We tested the utility and reliability of PharaohFUN by applying it to two case studies. These demonstrate its ability to infer the evolutionary trajectories and function of individual and entire sets of genes. The results support previous conclusions about evolutionary events related to terrestralization and central circadian clock proteins. For all these reasons, PharaohFUN is positioned as a tool with great potential for end-to-end phylogenetic studies in plant organisms, saving users large amounts of time and resources without results precision reduction.

## 1. Analysis pipeline

PharaohFUN is a web-based tool written in R, using the practical framework of the Shiny package. The complete analysis pipeline is summarized in Fig. 1A. PharaohFUN includes the proteomes of 167 species representing the main clades of the green lineage, namely Prasinodermophyta, Chlorophyta and Streptophyta (Sup. Table S1). As for the distribution of species by clades, PharaohFUN aims for a balanced representation avoiding clustering biases.

**Figure 1.**
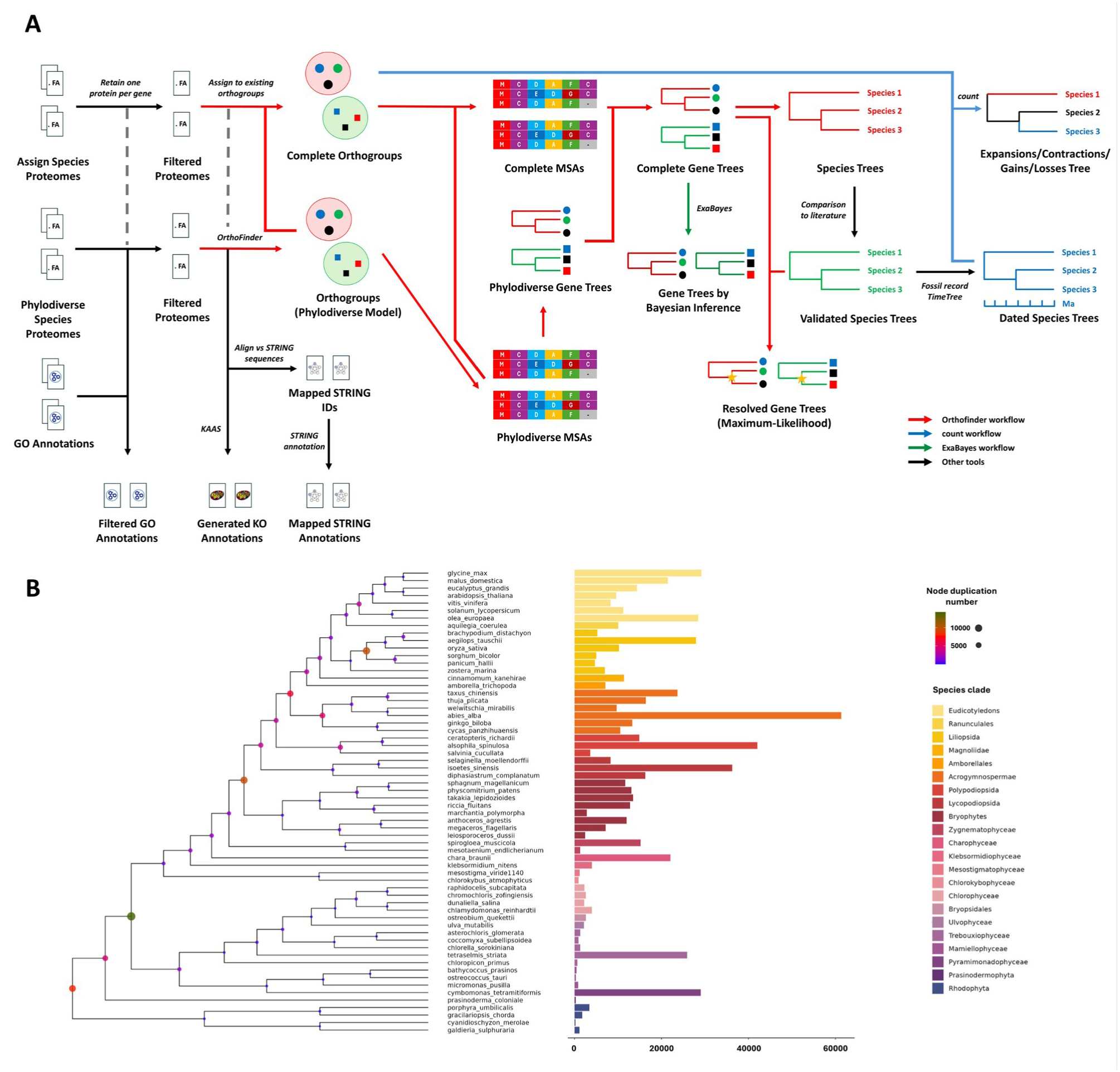
PharaohFUN’s model creation and composition. **A.** Analysis pipeline used for the generation of all input data used by PharaohFUN. Files derived from different software workflows are indicated through colored arrows. **B.** Phylodiverse species tree. **Left**: ultrameric species tree. The inferred gene duplication number of internal nodes is mapped by circle color and size. **Right**: gene duplication number of extant species. Clades are distinguished by different bar color according to legend.

This model has been generated using *Orthofinder* v3.0 (Emms and Kelly 2019; Emms et al. 2025) on the proteomes of the species mentioned in Sup. Table S1. The proteomes were previously filtered to include only the protein products of the primary transcript, as recommended in *Orthofinder* best practices. It ensures that the clustering remains unaltered by the appearance of multiple proteins with high similarity derived from the same gene. Therefore, we will hereafter refer to genes instead of proteins, since each protein sequence is representative of a single gene. The final model was generated in two steps. Firstly, a subset of 58 species homogeneously distributed throughout the complete Viridiplantae clade was selected, prioritizing species from different groups with refined annotation (Fig. 1B). The corresponding orthogroups (OGs) were calculated using the Multiple Sequence Alignment (MSA) function of the software, incorporating 4 rhodophyte species as outgroup to Viridiplantae within Archaeplastida. We will refer to this first model with a balanced sampling of species as the phylodiverse model. Then, we assigned the genes from the remaining proteomes to those precomputed OGs, adding them to MSAs and gene trees, and reconciling the latter with the bibliographically validated species tree (Strassert et al. 2021; Timilsena et al. 2022) (Sup. Fig. 1).

The final model contains 4,529,870 genes assigned to 97,610 OGs, representing 92.8% of the total genomes size. G50 statistic measures the number of genes in an OG such that 50% of the genes are in OGs of that size or larger. It corresponds to is 437, and 56 OGs contain genes from all species. Certain species exhibit a significantly higher number of identified gene duplications compared to other species within their respective clades (Fig. 1B). Notable cases are those of *Mesostigma viride*, *Spirogloea muscicola*, *Abies alba, Alsophila spinulosa* and *Cymbomonas tetramitiformis* attributed to whole-genome duplication (WGD) or triplication events and polyploidy (Cheng et al. 2019; Liang et al. 2020; Stull et al. 2021; Huang et al. 2022; Qiao et al. 2022; Cui et al. 2023; Gyaltshen et al. 2023).

Due to the variation in the number of genes per species in the OGs, an analysis of their contractions and expansions was performed using *count* (Csűös 2010) software with a fossil-calibrated species tree. The resulting tree from *Orthofinder* workflow was dated using the relaxed molecular clock model implemented in the R package *ape*. The selected model accounts for different rates of evolution among nodes drawn from a gamma distribution, with constraints derived from fossil record and TimeTree (Kumar et al. 2022) (Sup. Table S2). To allow for the reconstruction of ancestral states in all OGs, an asymmetric Wagner parsimony model was used. The software’s default gain penalty was relaxed to 4, to account for horizontal gene transfer (HGT) and whole genome duplications (WGDs) events known in this clade. (Wickell and Li 2020; Ma et al. 2022; Clark 2023).

Finally, existing Gene Ontology (GO) annotations for each proteome were downloaded and KEGG annotations were generated using KAAS (Moriya et al. 2007) with 20 photosynthetic eukaryotic organisms as reference. In the case of STRING data (Szklarczyk et al. 2019), the proteomes were mapped against the database sequences, using a 95% identity threshold, to associate the PharaohFUN IDs with those of STRING and generate the table of physical interactions. The type of evidence for the interaction was also associated: direct evidence or interolog.

## 2. Tool architecture

The OGs, gene trees, expansion/contraction/gain/loss models and annotations derived from the described workflow served as the basis of the results provided by PharaohFUN for each search. The complete architecture of the application is shown in Fig. 2.

**Figure 2.**
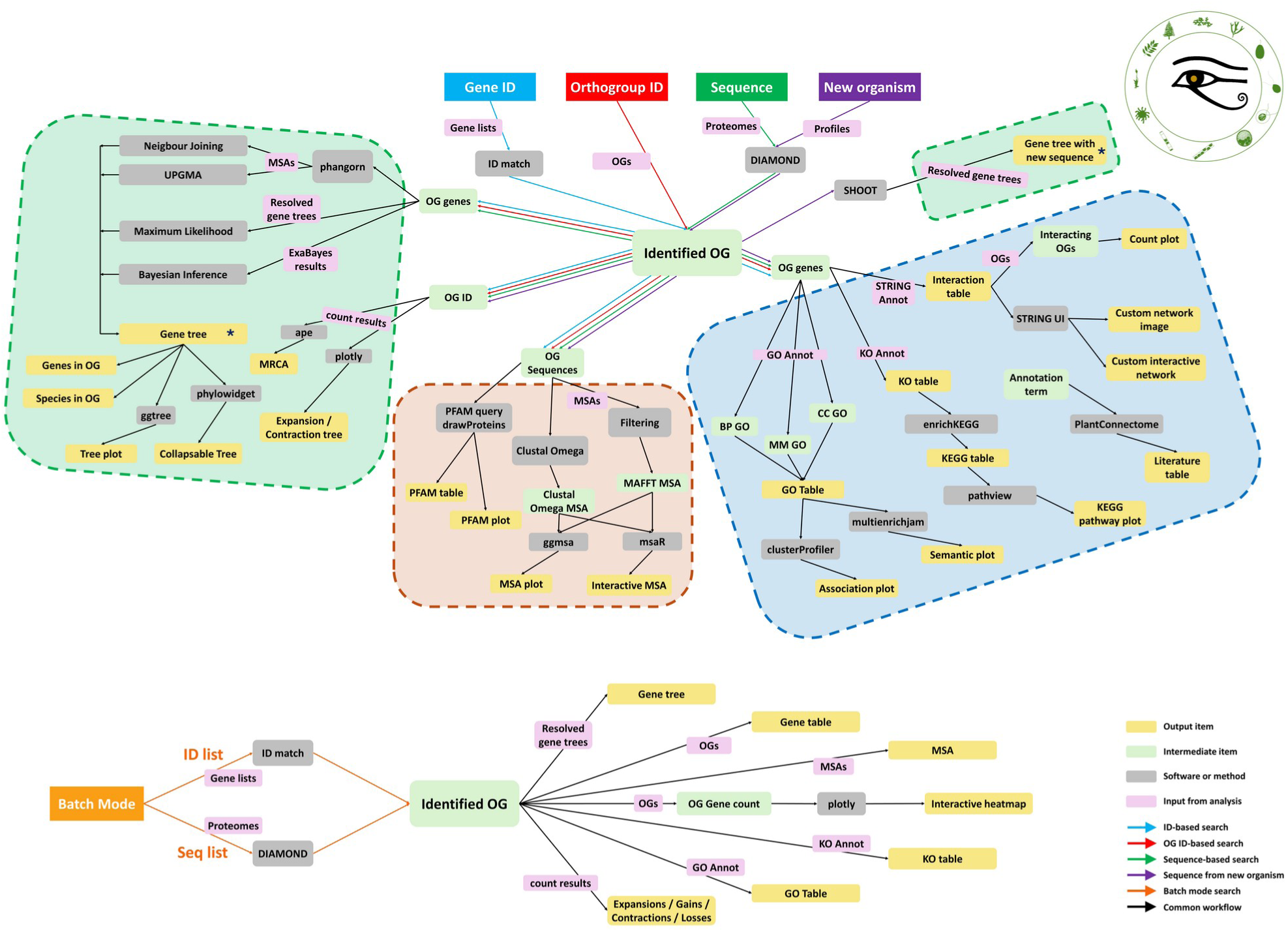
PharaohFUN architecture. The tool is divided into three main modules, with five different search modes. Each search mode is associated with a different colored arrow, and the common steps are represented in black. Color boxes stand for different types of objects, as determined in the legend. The asterisk marks that the elements are equivalent, and all outputs derived from “Gene tree” are also produced from “Gene tree with new sequence”.

The first step of any analysis with PharaohFUN consists of the identification of the OG of interest and its position in the tree of the input. The tool is designed to be flexible with respect to the input type to solve the ambiguity problems that exist in many databases. There are 5 search options: by gene ID (typically following the notation used in Phytozome (Goodstein et al. 2012) and Phycocosm (Grigoriev et al. 2021)), by sequence, by OG ID, in batch mode and by sequence of an organism not present in the database. In this last case, the DIAMOND aligner (Buchfink et al. 2015) is used to search for the most similar sequence in the supported proteomes when the input sequence actually belongs to a supported organism or to one evolutionarily close to some of the supported species. When the sequence belongs to a highly divergent organism from the ones supported, PharaohFUN incorporates a search based on the novel software tool SHOOT (Emms and Kelly 2022). This search mode identifies the most likely OG and includes the new sequence in the correct position of the gene tree. Finally, PharaohFUN incorporates a batch mode for the analysis of complete gene lists, allowing their inclusion as gene IDs or sequences from the same organism.

Once the OG to which the query belongs has been identified, PharaohFUN incorporates a series of features to extract information about its evolutionary context and function. They are organized in three main modules, as follows.

### 2.1. Gene Trees and Orthogroup Size Evolution

This module creates and represents gene trees corresponding to specific OGs, allowing for the selection of the species to be included and enabling users to restrict analyses to specific clades of interest. The tool incorporates four alternative methods for its construction: Maximum-Likelihood, Bayesian Inference, Neighbour Joining and UPGMA. While Maximum-Likelihood trees are directly derived from the analysis pipeline, the other methods rely on the MSAs calculated for each OG. Maximum-Likelihood corresponds to resolved gene trees derived from *Orthofinder* workflow, i.e. gene trees computed using *FastTree* and reconciled with the species tree (Emms and Kelly 2019). Neighbour Joining and UPGMA options directly compute the gene trees for the selected sequences from the corresponding MSAs, employing a bootstrap of 100, using the R package *phangorn* (Schliep 2011). Neighbour Joining resulting trees are then rooted by the midpoint of their longest branch. Trees determined by Bayesian Inference were precomputed using the Markov Chain Monte Carlo algorithm implemented in *ExaBayes* (Aberer et al. 2014). The generation of these trees was conducted using two independent runs and two coupled chains for each one, using a convergence criterion to stop the exploration of the probability landscape. The complete set of parameters employed is described in the file Sup. Table S3. Trees derived from all these building options are visualized using the *ggtree* package (Yu 2020). Their interactive exploration is made possible through *phylowidget* (https://github.com/sdwfrost/phylowidget). In addition, it is possible to collapse subclades to restrict the search, removing clades and not entire species.

The “Orthogroup Size Expansion/Contraction” tab complements the previous information exploring expansion, contraction, gain and loss events undergone by an OG throughout its evolution. This feature uses the output of *count* to explore changes in OG sizes mapped onto the species tree. It reconstructs the number of genes belonging to an OG in the common ancestors of each clade and the expansions, contractions, gains and losses suffered by the OG along each branch. In this tab, an interactive species tree is displayed with information on the number of genes within the OG in each node, along with the evolutionary events determined in each branch. Its MRCA is also highlighted.

### 2.2. Domains and Sequence Similarities

The focus of this module is to compare different sequences originating from the same gene tree. It assesses their potential functional relationships through the analyses of the PFAM domains they contain and the conserved and divergent regions according to an MSA. PharaohFUN implements a connection with the PFAM server to send a query for desired protein sequences. The results are presented as a structured table displaying the positions of the different domains in each sequence, along with information about them. Additionally, a graphical representation of the proteins representing domain locations is presented.

In the MSA tab, two methods are provided: ClustalOmega and MAFFT. The former uses the ClustalOmega algorithm (Sievers et al. 2011) to align selected sequences. The latter uses the complete OG MSA generated using MAFFT and filters out non-selected proteins. Columns that correspond only to gaps in the sequence subset are removed. This feature is useful when studying a specific clade of the gene tree, enabling the examination of sequence conservation within that group rather than in the entire tree. The output of this section consists of an interactive alignment viewer that can also be used to find motifs using regex notation.

### 2.3. Functional Annotation

One of the objectives of PharaohFUN is to facilitate the formulation of hypotheses about the function of proteins with deficient functional annotation leveraging their evolutionary relationship with proteins of known function. To achieve this, PharaohFUN implements four different analyses: GO enrichment, KEGG orthology, STRING interactions and literature-based annotation.

In the “GO Terms” tab, the gene ontology terms of a subset of genes in the tree are listed. This analysis functionally characterizes genes based on existing functional annotations of phylogenetically close genes A list of existing GO terms is displayed for each gene summarized in two different graphical representations. The first one is an emapplot visualizing relationships between different groups of terms based on connections by genes involved in different processes. The second representation is a treeplot clustering GO terms according to their semantic similarity, providing a summarized view.

In the “KEGG Orthology” (KO) tab, once genes are selected, their corresponding KO terms are identified. Subsequently, KEGG pathways are determined based on KO terms present in the selection. This analysis reveals the different molecular pathways in which the selected genes could be involved. The output is presented in a tabular format, and users can also choose to visualize a specific pathway to explore in which steps the selected genes act.

The “STRING Interactions” tab incorporates information on protein-protein interactions contained in the STRING database (Szklarczyk et al. 2019). The tool displays all the proteins in the organism that interact with each of the chosen ones, as well as information on the evidence supporting this interaction. In turn, the tool analyzes OG co-evolution by identifying to which OG each of the target proteins belongs to. Moreover, it also generates a graphical representation of the interaction network of a subset of proteins of interest and a link to the corresponding interactive network. For the proteins in the network, 3D structure and SMART domains information can be accessed.

Finally, the “Literature Annotation” tab provides a connection to the PlantConnectome web tool (Lim et al. 2024). This feature allows users to retrieve extensive information from plant science literature, obtained through the application of Large Language Models to process articles related to this field. Users can associate the results from the previous tabs with other bibliographically supported biological information. Additionally, a link to the paper from which each piece of information has been inferred is provided to expand or verify the accuracy of the results.

## 3. Case studies

PharaohFUN has been developed for two distinct types of studies. On the one hand, it allows the fine-grained study of the evolution of a gene of interest and the annotation of associated genes. On the other hand, it can simplify and accelerate phylogenomic studies for large numbers of genes. Its use in these scenarios is presented by two representative case studies.

### 3.1. Case study 1: Characterization of a putative CCA1 in *Helianthus annuus*

To demonstrate PharaohFUN’s utility for fine-grained studies, the sequence of a putative CIRCADIAN CLOCK ASSOCIATED 1 (CCA1) protein from *Helianthus annuus* (entry A0A251TYD1) was downloaded from UniProt and analyzed using the "Sequence from new organism" function. CIRCADIAN CLOCK ASSOCIATED 1 (CCA1) is a central regulator of circadian rhythms largely conserved throughout Viridiplantae. *A. thaliana, B. rapa, A. hypocondriacus, M. polymorpha, O. tauri* and *P. patens* were selected as species of interest. The resulting gene tree showed a division into three main clades (Sup. Fig. 2). The query protein was located in that shown in Fig. 3A, split in two subclades.

**Figure 3.**
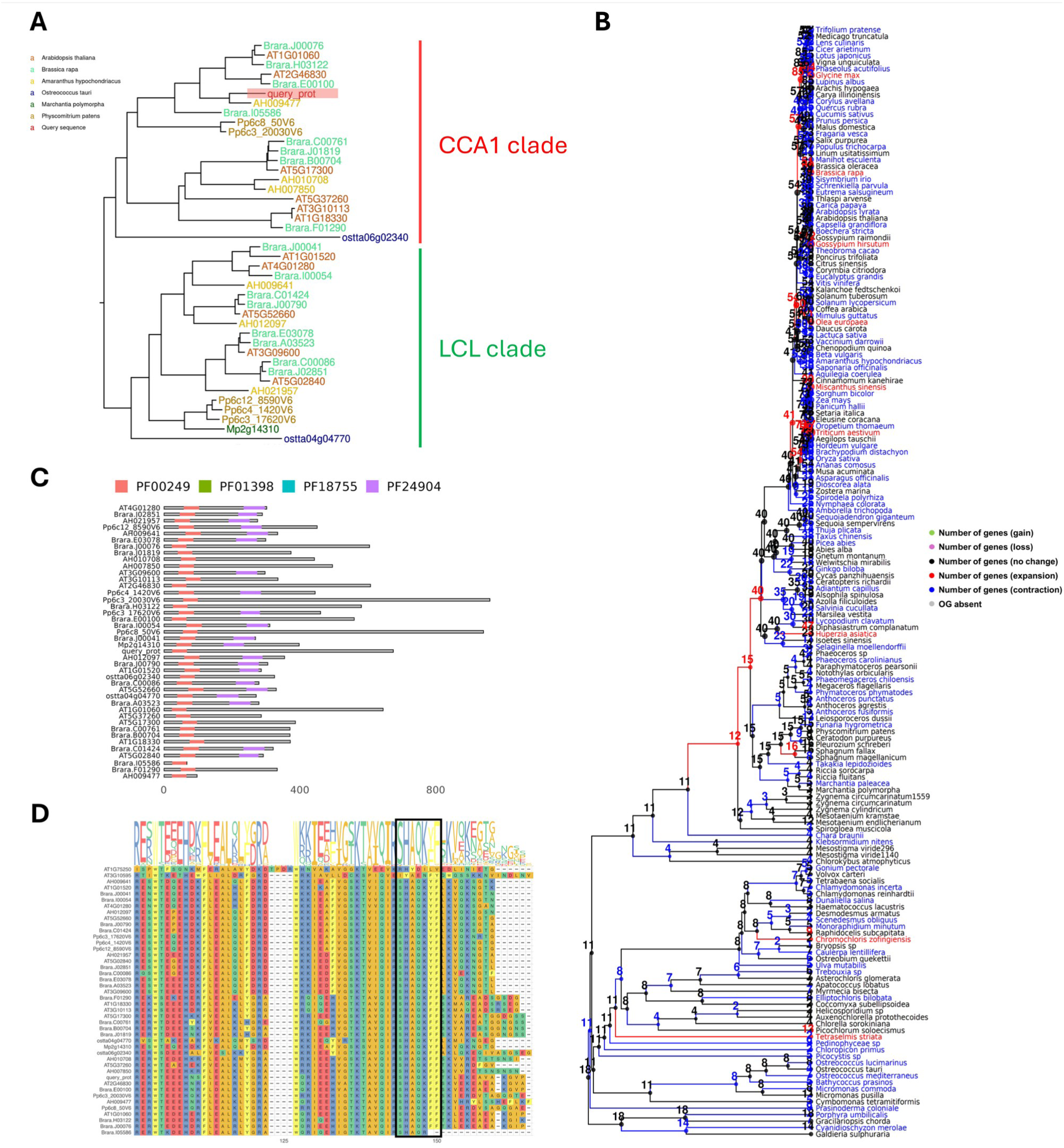
Overview of results for *Helianthus annus* putative CCA1 evolution using PharaohFUN. **A.** Gene tree for *Helianthus annus* putative CCA1 orthogroup using *A. thaliana, B. rapa, A. hypocondriacus, M. polymorpha, O. tauri* and *P. patens* as reference species. The query gene is highlighted in red. **B.** Size evolution of orthogroup through species tree. **C.** PFAM Domains location in proteins corresponding to CCA1/LCL clade. **D.** MSA in the vicinity of the conserved SHAQKYF domain, highlighted by a black box, for the protein sequences in CCA1/LCL clade plus two additional *A. thaliana* proteins from the same orthogroup.

The first subclade contained *A. thaliana* CCA1 and LHY, along with the REVEILLE family members RVE1, 2 and 7/7-like. It also included the previously identified CCA1 orthologs in *P. patens* (CCA1a and CCA1b) and *O. tauri* (Okada et al. 2009; A.-M. Linde et al. 2017). In agreement with this clade representing CCA1/LHY orthologs, no *M. polymorpha* sequences were included (A.-M. Linde et al. 2017). We can observe that the protein of *H. annus* groups with that of *A. hypocondriacus*, also from superasterids clade. As for *B. rapa*, it presents the gene distribution indicated in (Lou et al. 2012) as a consequence of a whole genome triplication and gene loss after its divergence from *A. thaliana*. Thus, two LHY copies and one CCA1 are present, together with a higher number of RVEs. However, it presents an additional protein not described in the previous study. This protein is Brara.I05586, present in the LHY/CCA1 clade with lower sequence similarity to these two than the aforementioned copies.

The second clade was constituted by proteins traditionally referred to as the LHY-CCA1-Like (LCL) family. Within this clade, a gene from *M. polymorpha* known as MpRVE, and one from *O. tauri* were identified, supporting predictions from previous analysis (A.-M. Linde et al. 2017). Similarly, the known increase in the number of copies of RVE 4/8/6 and the retention of a single copy of RVE3 and RVE5 in *B. rapa* (Lou et al. 2012), as well as the existence of three PpRVE copies (A.M. Linde et al. 2017) were detected.

To further explore the outcomes derived from the gene tree, we used the rest of functionalities in PharaohFUN. Notably, the analysis of orthogroup contraction/expansion revealed (Fig. 3B) a contraction in the branch leading to *M. polymorpha* and expansions in that of *B. rapa*. To gather more evidence and refine previous results, a PFAM domain analysis was conducted with PharaohFUN. The proteins from the gene tree were found to contain the Myb-like DNA-binding domain (PF00249) and often a REVEILLE 6-like domain (PF24904). This domain is known to be highly prevalent in transcription factors that constitute the central circadian clock (Carré and Kim 2002). Accordingly, *Helianthus annus* putative CCA1 ortholog showed PF00249 domain. The rest of the orthogroup proteins present this domain in different locations and combinations (Sup. Fig S3). Next, we performed MSA using the sequences of these proteins plus two different sequences from an external clade (AT1G75250 and AT3G10595). Using the PharaohFUN function “Find motifs”, the conserved SHAQKYF motif was searched for. This motif is typically associated with RVE and CCA1 orthologs (A.-M. Linde et al. 2017). It was identified in all proteins (or its variant SHAQKFF) except for the two Myb-like DNA-binding proteins not clustered in the CCA1-LCL clade (Fig. 3D). Interestingly, in the *B. rapa* Brara.I05586 protein, this domain is truncated, which may have led to loss of function and explain its distance with respect to both CCA1 and LHY.

We applied the remaining functionalities in PharaohFUN to leverage the evolutionary results obtained thus to elucidate potential functions for the uncharacterized gene in this example. It is important to consider that the species whose genes are used to transfer functional information can greatly impact the accuracy of the results. Therefore, the study of clades in the tree is crucial for selecting genes that may carry information of interest for a particular study gene. Exploration of GO terms from genes belonging to the CCA1 clade revealed processes mostly related to circadian rhythm transcription and response to cold and, with lower abundance, auxin biosynthesis, seed germination, flower development, catabolic processes, response to blue light and temperature compensation of the circadian clock (Fig. 4A). In addition, the KO annotation confirmed the presence of homologs of CCA1 and LHY in the orthogroup, highlighting their role in pathways related to circadian rhythm (Fig. 4B). The literature annotation feature provided valuable biological information, including descriptions of relationships such as complex formation with LHY (which belongs to the same OG), repression of TOC1 through binding, control over circadian rhythms, and more than 200 further relationships (Fig. 4C). These annotations can be used to experimentally test hypotheses regarding conservation of function among predicted orthologs.

**Figure 4.**
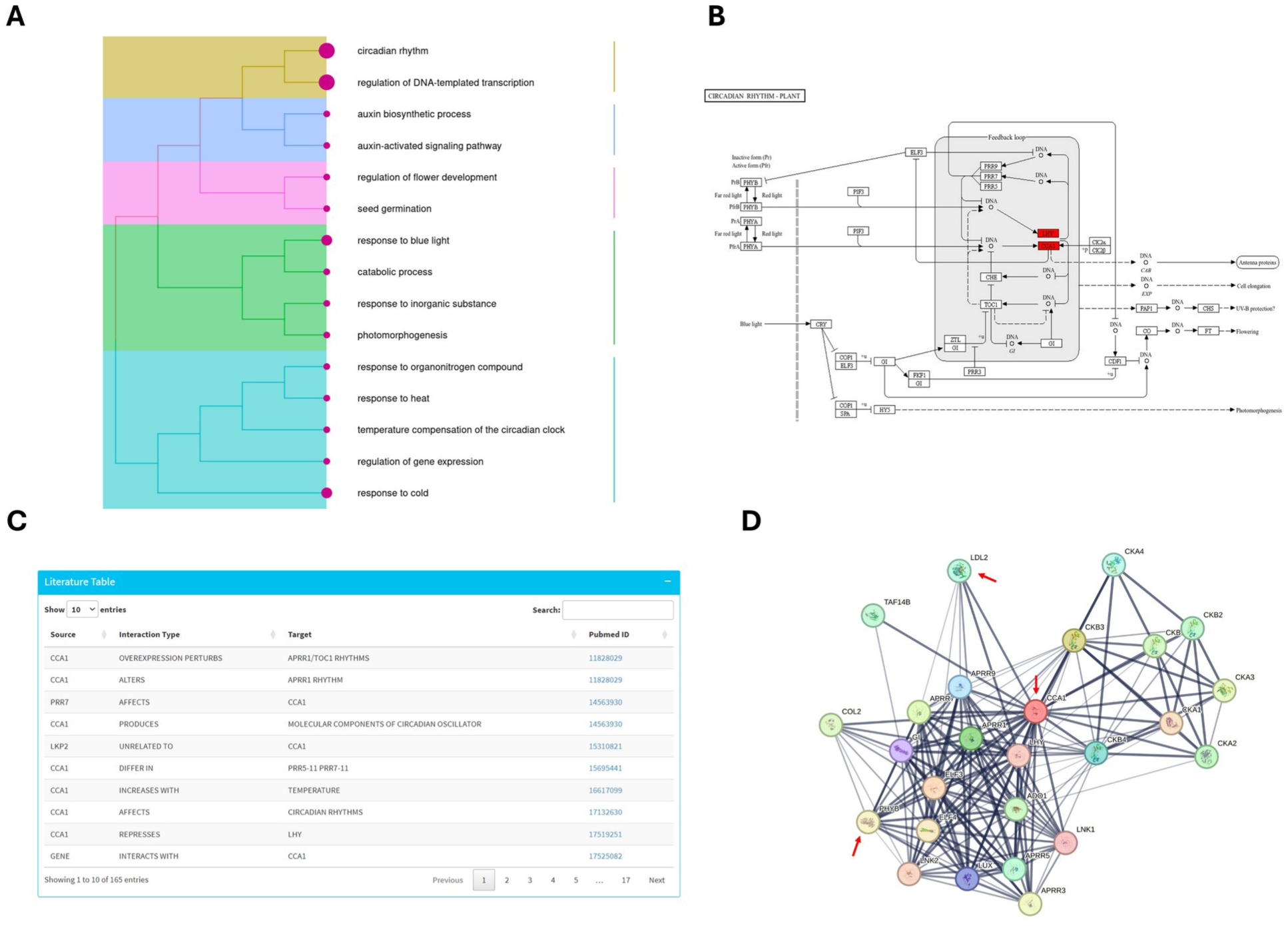
Overview of results for CCA1-clade functional annotation using PharaohFUN. **A.** Treeplot showing clustered GO terms for biological processes. **B.** KEGG pathway for Circadian Rhythms in Plants, highlighting genes in the clade (CCA1 and LHY) in red. **C.** Example of some rows of literature annotation result for “CCA1” using exact search. **D.** STRING interaction network of AT2G46830 gene. Red arrows indicate genes from orthogroups that appear in the first rows of conserved interactions results.

**Figure 5.**
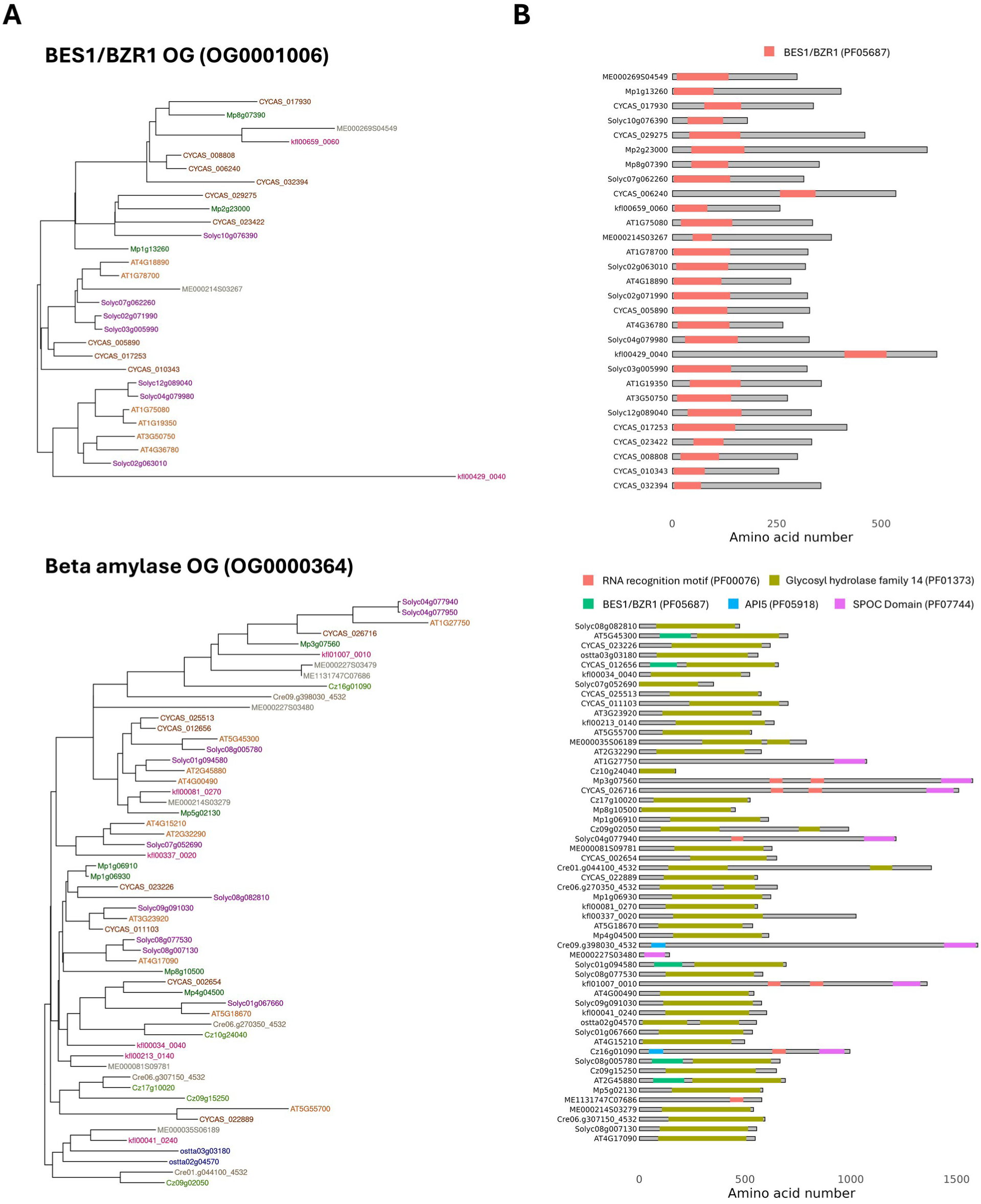
Results for the different OGs associated to BES1 family. **A.** Gene trees. **B**. PFAM domains location. Panels: **Up** OG corresponding to BES1/BZR1 TFs, **Down** OG corresponding to beta amylases.

Finally, to study coevolution between OGs with interacting proteins, the STRING function was employed, selecting as input the CCA1 orthologs identified in *O. tauri*, *P. patens* and *A. thaliana*. The OG whose proteins interacted with this subset a higher number of times was found to correspond to ID OG0001357 (Sup. Table S4), which was entered into the OG search mode developed for this purpose. This revealed an orthogroup comprising PHYTOCHOME A-E. The role of the interaction between CCA1/LHY and PHYB in the response to far-red light has been well described, regulating processes such as shade avoidance, circadian clock entrainment and photomorphogenesis (Yeom et al. 2014; S. Wang et al. 2022; Bian et al. 2025). These proteins also appeared when plotting the interaction network of CCA1 in PharaohFUN (Fig. 4D). The second one, OG0000363, was found to contain a cluster of genes related to Lysine-Specific Demethylase 1 (LSD1)-like histone demethylases LDL1/2. In addition to appearing in CCA1 interaction network (Fig. 4D), there is evidence in the literature that LDL1/2 can interact with CCA1 and LHY to repress TOC1 expression (Hung et al. 2018) and that these proteins are conserved in the green lineage (Martignago et al. 2019), validating our results.

This example showcases the strengths of PharaohFUN, which arise from its ability to combine multiple techniques for phylogenetic analysis, gene trees, PFAM domains and MSA to identify potential orthologs and study their evolutionary history, even when studying species not included in the dataset. Based on these analyses, functional annotations can be associated with unknown proteins (Gil et al. 2023).

### 3.2. Case study 2: Analysis of the evolutionary history of the complete endowment of TFs of *A. thaliana* across Viridiplantae

PharaohFUN can also be used as a tool for large-scale phylogenomic studies of any gene set of interest. To demonstrate this, an analysis was performed to explore the evolutionary history throughout Viridiplantae of the complete set of transcription factors (TFs) of *Arabidopsis thaliana*. Traditionally, this history has been generated individually for each TF family or even their subclades. Here we show how PharaohFUN can be used to perform a global study in a quick and easy way. Our results are shown to be consistent with the literature as demonstrated in terms of tree topologies and ancestral state of OGs. The orthologs recovery by PharaohFUN as a function of evolutionary distance has been also tested.

#### 3.2.1. PharaohFUN reconstructs known orthology and gene tree topologies

The complete set of transcription factors sequences from *A. thaliana* was inputted to PharaohFUN Batch Mode selecting 54 species in the green lineage. Using the classification of PlantTFDB (Jin et al. 2017), the 51,892 sequences resulting from this search were classified into 58 families of TFs. PharaohFUN Batch Mode results were used to analyze the OGs that composed each family, their MRCA, their associated number of genes per species and their PFAM domains (Sup. Table S5-S6). Individual OGs were identified for 37 of the 58 families, while 10 were split into more than 5 OGs (Sup. Table S5). 8 OGs belonged to more than one family. This division into several OGs for specific TF families followed two main reasons: different MRCAs for subgroups of the same family and modification of functional domains in subgroups. Accordingly, many of them reflect accurately the evolutionary history of TF families and the true topologies of their gene trees. Examples for this are BES1, WRKY and C2H2 families.

The BES1 TF family is an example of a domain-based division into different OGs. PharaohFUN split the BES1 family into two OG (Fig. 4A). The first one of them was present throughout the entire Viridiplantae from its MRCA and presented a Glycosyl hydrolase family 14 domain as well as some N-terminal BES1/BZR1 domains (Fig. 4B). The MRCA of the second OG arose in the ancestor of the clade constituted by Klebsormidiophyceae+Phragmoplastophyta, and only possessed the BES1/BZR1 domain. Genes from chlorophyte algae in the first OG revealed only the hydrolase domain, whereas BES1/BZR1 did not appear. Therefore, one OG of this family reflects the evolution of BES1 TF and the other represents the evolution of beta amylases that at a particular point acquired BES1 domains. This result is in agreement with previously published work indicating that streptophyte algae and land plants possess this TF family, unlike chlorophytes (Wilhelmsson et al. 2017; Mecchia et al. 2021).

The PharaohFUN results regarding the WRKY family, one of the largest TF families in *A. thaliana*, were analyzed to test its performance on families with complex evolutionary histories. The WRKY family was divided into 3 OGs. The first one corresponded to WRKY TFs (combining groups I, II and III) and the second, to the TIR-NB-LRR group (Kalde et al. 2003; Wang et al. 2015; Villacastin et al. 2021). As for the third OG, it was found to correspond to MAPK/ERK kinase kinases, with only one protein (WRKY19/MEKK4) exhibiting a WRKY domain in *A. thaliana*, in addition to its kinase domain. In turn, the topology of the WRKY tree reflected the phylogenetic studies performed on this group. Group III maintained the organization in four clades described in (Wang et al. 2015), while group I appeared independently and II showed the separation of the clades II-a, II-b, II-c, II-d and II-e (Huang et al. 2012) (Sup. Fig. S4A). Group II did not constitute a single entity, but subgroups IIa/b, IId/e y IIc were confined to phylogenetically distinct groups. This organization was already pointed out in (Mohanta et al. 2016) as a consequence of performing a phylogeny based on a larger number of plant species instead of just dicots. Thus, our results are consistent with their revisions on this family phylogeny. In the same study, the authors pointed out the existence of chimeric proteins with a WRKY and a TIR, LRR or kinase domain occurring mostly in flowering plants and not present in all plant species, fully explaining the division into orthogroups returned by PharaohFUN. Finally, as for the TIR-NB-LRR OG, the analysis of its evolutionary history showed a complex scenario, with an origin prior to the division between chlorophytes and streptophytes, but abundant losses in multiple lineages (Sup. Fig. S4B). Among them, monocots stand out, while some chlorophyte algae such as *Chromochloris zofingiensis* and streptophytes such as *Klebsormidium nitens* exhibited representatives of these proteins, in agreement to previous findings (Tarr and Alexander 2009; Andolfo et al. 2019; Shao et al. 2019). Traditionally, this family was believed to have appeared in streptophyte algae due to the low number of representatives of the different clades of Chlorophyta included in the corresponding analysis (Shao et al. 2019), which exemplifies the importance of the inclusion of these organisms in evolutionary studies and justifies the high number of them included in our tool.

Finally, C2H2 is the family that was split into the largest number of orthogroups, 18. In this case, this division was not artifactual either, but faithfully represented the discordance presented in (Li et al. 2022) between the phylogeny of C2H2 zinc-finger proteins (C2H2 ZFPs) constructed for multiple plant species and the sequence similarity of the ZFPs in the individual species. Thus, the mapping of the different OGs onto the AtZFPs tree reported in that study allowed us to observe how the three main OGs corresponded to the three largest contiguous clades with proteins of the same global group (groups III, IVb and IVc) in the AtZFPs tree, whereas the appearance of proteins of different groups in the subclades promoted the split of the corresponding proteins into different OGs (Sup. Fig. S5).

This demonstrates that PharaohFUN’s OGs reflect the evolutionary trajectories of the gene families and that they coincide with reported tree topologies. Thus, it can be reliably used for orthology and phylogenetic studies.

#### 3.2.2. PharaohFUN correctly traces the evolutionary history of OGs

Our tool goes beyond gene trees and allows users to explore the evolutionary history of the different OGs, in terms of the distribution of the number of genes per species identifying expansion/contraction/gain/loss events. In addition to the individualized study of the aforementioned TF families, we used PharaohFUN to perform a global and integrated analysis of all these genes. Accordingly, the number of orthologs of *A. thaliana* genes was determined for each TF family over Viridiplantae. Fig. 6A shows the distribution of genes in each family, consisting of the sum of genes from all OGs associated with each family. For the same representation separated into OGs instead of TF families (direct PharaohFUN output), refer to Sup. Fig. S6. Overall, there was a substantial number of TF families without representatives in chlorophyte microalgae, whereas those that were present exhibited a lower number of genes compared to vascular plants. Likewise, streptophyte algae were postulated as a middle ground between both groups. Although most TF families are present in all or some organisms of this paraphyletic group, their gene abundance was low, similar to those of chlorophytes.

**Figure 6.**
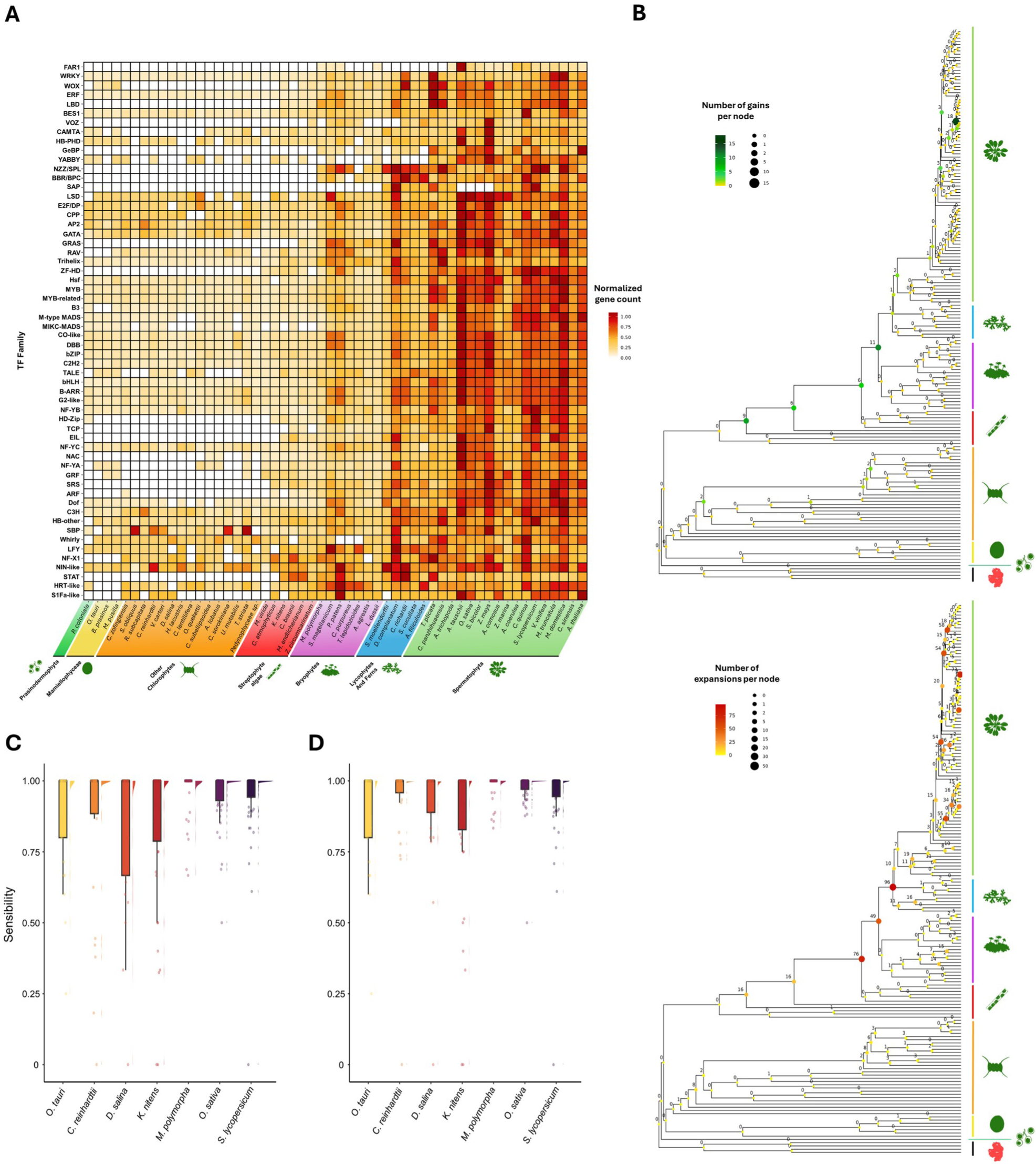
Evolution of the TF families from *A. thaliana* in Viridiplantae. **A.** Heatmap showing the normalized gene number per TFs family in 54 species supported by PharaohFUN. TF families are ordered based on their abundance pattern similarity through the different organisms. **B.** Species tree with the number of gene acquisition events mapped onto its internal nodes. Up: OG gains. Down: OG expansions. **D.** Raincloud plot showing the distribution of sensitivity (recovery rate) in the identification of genes of each TF family in each test species using *A. thaliana* genes as a source. **E.** Raincloud plot showing the distribution of sensitivity in the identification of genes of each TF family in each test species using *A. thaliana*, *B. prasinos*, *V. carteri*, *P. patens* and *S. bicolor* genes as source. The raincloud plot depicts a typical boxplot along with the individual jittered measure points and their density is represented at the side of the boxplot.

These results suggest that an important number of TF families exhibited today by *A. thaliana* arose prior to land colonization by the green lineage possibly promoting this key event in life evolution. To further explore this, we used the expansion, contraction, gain and loss events in OGs through the entire species tree identified using the Batch mode in PharaohFUN. This analysis also identified the position of the MRCA associated with each OG (Sup. Fig. S7). In addition, gains at the level of each internal node were calculated (Fig. 6B and Sup. Fig. S8). Our results indicated that many OGs corresponding to *A. thaliana* TF families are not of recent evolutionary origin, instead a large number (83) was already present in Viridiplantae MRCA. Likewise, the number of OGs that appeared in the MRCAs of streptophyte algae, Embryophyta and Magnoliopsida is particularly relevant, pointing to a progressive increase in the repertoire of TFs. Also interesting is the high number of Arabidopsis-specific gains (18), associated to OGs present only in this genus. OGs originated between the emergence of Streptophyta and Embryophyta were related to the TF families shown in Table 1. It should be noted that no OG present in *A. thaliana* could have emerged in a clade that does not contain this species. This explains the absence of new OGs in the common ancestor of other groups. MRCA of TF families was compared to that determined by (Wilhelmsson et al. 2017) for families that allowed direct comparison and to (Jin et al. 2017) for the rest. To allow comparison, the MRCA of the oldest OG for each TF was manually curated, performing a domain analysis to discard OGs that did not correspond to the TF family, as we exemplified for BES1.

**Table 1.** TF families associated to the OGs that emerged in different ancestors of *A. thaliana*.

| Node | TF family emergences |
| --- | --- |
| Root (Archaeplastida) | ARR-B / bHLH / bZIP / C2H2 / C3H / CO-like / CPP / DBB /<br>E2F_DP / G2-like / GATA / HB-other / HSF / M-type MADS /<br>MYB / MYB-related / NF-X1 / NF-YA / NF-YB / NF-YC / NIN-like / TALE / WRKY / YABBY |
| Viridiplantae | AP2 / B3 / Dof / ERF / SBP / Trihelix |
| Chlorophyta + Streptophyta | CAMTA / HD-zip / S1Fa-like / Whirly / WOX |
| Streptophyta | ARF / GRF / MICK MADS / RAV / SRS / ZF-HD |
| Phragmoplastophyta +<br>Klebsormidiophyceae | BES1 / EIL / LBD / LFY / LSD / NAC / STAT |
| Phragmoplastophyta | HRT-like / TCP |
| Anydrophyta | BBR-BPC / GRAS / VOZ |
| Embryophyta | FAR1 / GeBP / NZZ_SPL |
| Tracheophyta | SAP |

Thus, our inference matched theirs for 55 out of 58 families. As for the remaining three, we placed CAMTA MRCA in Chlorophyta+Streptophyta, consistent with (Jin et al. 2017) but prior to the inference in (Wilhelmsson et al. 2017), which may be due to the inclusion of more chlorophyte algal species in our dataset. In the case of VOZ, we detected its origin in Anydrophyta (Rensing 2020), because of its presence in *Zygnema circumcarinatum*, *Zygnema cf. cylindricum* and *Mesotaenium kramstae* genomes, which were not included in their datasets. Finally, we inferred the emergence of LSD in Phragmoplastophyta+Klebsormidiophyceae instead of Viridiplantae. Unlike the other studies, the aim of this one is to show the recoverability of orthogroups based only on *A. thaliana* proteins, so there may be other OGs that contain proteins with the domains corresponding to LSD TFs in chlorophytes but are the result of other acquisitions without conservation of the rest of the sequence, resulting in the observed results. Emergence and gain of the total OG set depicted in Fig. 6B, including those that did not constitute the main group of the corresponding TF family are listed in Sup. Table S7.

In addition to the emergence of new OGs and TF families, the number of genes associated to them has changed significantly during the evolution in Viridiplantae. By studying the number of expanded OGs in the MRCA of the different clades, we identified four key events in the development of *A. thaliana* TF families. Namely, the common ancestors of Anydrophyta, Embryophyta, Tracheophyta and Magnoliopsida (Fig. 6C and Sup. Fig. S9). In the case of Embryophyta, it correlates with new TF-associated OGs emergence events as described above. However, the MRCA of Anydrophyta exhibited one of the most notable duplication events without the emergence of many OGs, second only to Tracheophyta.

OGs contractions in TFs families were also characterized (Sup. Fig. S10). Ancestors internal to Bryophyta, Lycopodiopsida and Polypodiopsida clades exhibited important contractions. Such events were more frequent at tips, probably reflecting a current evolutionary exploration, not necessarily successful adaptations that would be transmitted in the future. As a special case of these, losses (contractions leading to gene absence in the corresponding species) were especially relevant in chlorophytes (Sup. Fig. S11).

Many studies have been conducted to shed light on the molecular adaptations that took place in the aquatic ancestors of land plants that promoted their terrestrialization. With this case study, we show the utility of PharaohFUN to conduct this type of global analyses, saving time and making them available for every user. We have shown that the emergence of new OGs comprising TFs in *A. thaliana* has played an important role since the emergence of Streptophyta, accumulating new families that plants would later exploit for terrestralization. These results are in agreement with published work that links these innovations, for example, with high light irradiance tolerance in streptophyte algae (Rensing 2020). Some of the species from this group proliferate in shallow water or seasonal pools, requiring pre-adaptations to terrestrial conditions and, consequently, regulators of gene expression (Wilhelmsson et al. 2017; Lai et al. 2020). These results are also in agreement with the conclusions of other works which identified these OG gains to have taken place before the conquest of land (Wilhelmsson et al. 2017; Bowles et al. 2024). The specific OG expansions identified by PharaohFUN showed that this phenomenon is prevalent in the ancestors of Anydrophyta and the different Embryophyta clades, enabling the co-option of the existing families for other purposes (Table 1 and Sup. Table S7). These expansions have been thoroughly studied, with studies supporting that they allowed the preassembly of signaling and regulatory pathways that would be later used by land plants (Wilhelmsson et al. 2017; de Vries et al. 2018; Rensing 2020; Serrano-Pérez et al. 2022; Hernández-García et al. 2024). Originally, their function was different from that exhibited by this group, for example, regarding phytohormone signaling (de Vries et al. 2018; Serrano-Pérez et al. 2022). Regarding contractions, although they were prevalent in certain clades of Viridiplantae, their influence on shaping the *A. thaliana* TFs toolbox was lower than that of expansions and new OGs emergence. This is further supported by the literature on contractions of TFs families being determinant in clades such as Bryophyta, but exhibiting a reduced number for the branches leading to *A. thaliana* (Wilhelmsson et al. 2017).

Taken together, these results illustrate how PharaohFUN can accurately trace globally the evolutionary history of the different OGs, assisting the formulation of hypotheses about the evolution of protein families of interest based on the changes in OG size. Ancestral State Reconstruction and expansion/contraction studies are widely used to hypothesize how gene gains and losses influenced the evolutionary trajectories of extant organisms (Gàlvez-Morante et al. 2024). However, their performance requires highly specific training. PharaohFUN democratizes access to these studies for photosynthetic organisms, with the accuracy shown in these case studies. It highlights the ability of PharaohFUN to perform not only individual fine-grained studies but also global phylogenetic ones.

#### 3.2.3. PharaohFUN enables the recovery of orthologs at high evolutionary distances

In this case study, only the evolutionary history of OGs associated with *A. thaliana* TFs was traced. As a result, OGs not present in this species were not included in the analysis. We have determined the abundance of these to measure PharaohFUN’s ability to retrieve genes with a shared evolutionary history. Accordingly, we downloaded the complete TFs sets in 7 different species from PlantTFDB and assessed how many appeared in our results to evaluate the sensitivity of our tool as a function of divergence time between the source species and the target species (Sup. Table S8). The result was a mean sensitivity of 0.84 across all species, with lowest value corresponding to *K. nitens* (0.74) and highest to *O. sativa* and *S. lycopersicum* (0.93). At the level of individual TF families (Fig. 6E), the median of their sensitivities was 1 for all species (i.e., most families of each species were fully recovered by orthology with *A. thaliana*). The family with the lowest recovery rate in the different species was C2H2, with a median of 0.44. The overall high percentages point to a low abundance in other species of genes in OGs associated with TF families not present in *A. thaliana*. Furthermore, the distribution of sensitivities along the species tree suggests high effectiveness in identification of orthologs even at large evolutionary distances, probably due to the homogeneous species sampling employed in PharaohFUN.

Subsequently, TFs from *B. prasinos*, *P. patens*, *S. bicolor* and *V. carteri* were used as additional input to that of *A. thaliana* and the recovery rates of the previous seven species were recalculated (Fig. 6F and Sup. Table S9). By introducing representatives of clades in which *A. thaliana* is not present, OGs whose origin can be traced back to the common ancestor of those clades could be identified, improving sensitivity despite the very small number of new species added. The overall mean sensitivity was 0.90 in this case. The species that now presented the highest percentage of TFs recovered was *O. sativa* (0.97), probably due to the influence of *S. bicolor* in the identification of TFs originated at Liliopsida MRCA. On the other hand, the species with the lowest value continued to be *K. nitens*, unsurprising because species encompassing common ancestors of this organism not shared with *A. thaliana* were not added. The same phenomenon is detected in *S. lycopersicum*, which shows a minimal recovery rate improvement. The median recovery rate per TF family for each species remained 1 without exception, and the family with the lowest value was LFY (0.75). C2H2 family now showed a median of 0.88, compared to 0.44 without the input of the new species. This demonstrated that new OGs were recovered by improving the sampling in clades, not related to a poor performance of the tool, but to true evolutionary histories of different OGs.

The previous results demonstrate how PharaohFUN is able to recover a large percentage of the proteins associated with TFs in *A. thaliana* from other plant species, while maintaining a correct topology of the resulting gene trees and an overall correct MRCA inference of each group. We have demonstrated its adequacy with results from well-established tools for TFs analysis such as TAPscan (Petroll et al. 2025). TAPscan focuses on transcription-associated proteins and excels in this particular case study, due to being domain-oriented software and leveraging a balanced species sampling, using a similar two-step model creation as that of PharaohFUN. This domain orientation is especially advantageous for TF evolution studies, in which proteins are characterized by extensive functional domains that can be used to classify each family. It avoids the fragmentation into OGs that can be observed for some large families with widespread domains across the plant genomes. However, PharaohFUN does not use this paradigm since it may not be valid in the case of proteins that do not present functional domains and exhibits important limitations when applied to proteins containing domains of reduced extension. In these instances, the phylogenetic signal is not sufficient to create a reliable phylogeny of the proteins under study (Sjölander et al. 2011). PharaohFUN, being a general tool for the study of any protein set, is OGs-oriented like other tools of this nature, such as eggNOG-DB (Hernández-Plaza et al. 2023) and pico-PLAZA (Vandepoele et al. 2013). Therefore, the presented case study is one of the most potentially challenging analysis that can be carried out by PharaohFUN when compared against domain-oriented software, due to the large family sizes being evaluated. However, we have been able to verify how a high number of families (37) were contained in a single OG, how the expected split into OGs is usually motivated by the evolutionary history of the family as corroborated by specific literature, and how the evolutionary events at the species level were generally in agreement with the results from state-of-the-art specialized software such as TAPscan.

All these results demonstrate the robustness of PharaohFUN as a tool to detect orthologs of genes of interest at large scale. The selected species set allow for orthology relationships to be detected at high evolutionary distances, by combining uniform sampling of clades and the inclusion of streptophyte algae. This case study showcases the great amount of evolutionary data that PharaohFUN outputs by using a simple gene or sequence list as input.

## 4. Comparison with other web-tools and databases

Currently, there is a limited collection of tools available for quickly identifying orthologs of plant genes, and few of them include a diverse range of plant species in their databases. Notable exceptions are SHOOT (Emms and Kelly 2022), FunTree (Sillitoe and Furnham 2016), eggNOG-DB (Hernández-Plaza et al. 2023), PhylomeDB (Fuentes et al. 2022) or, exclusively for plant species, pico-PLAZA (Vandepoele et al. 2013), which cater to specific objectives and capabilities differing from those of PharaohFUN, as summarized in Table 2.

**Table 2.**
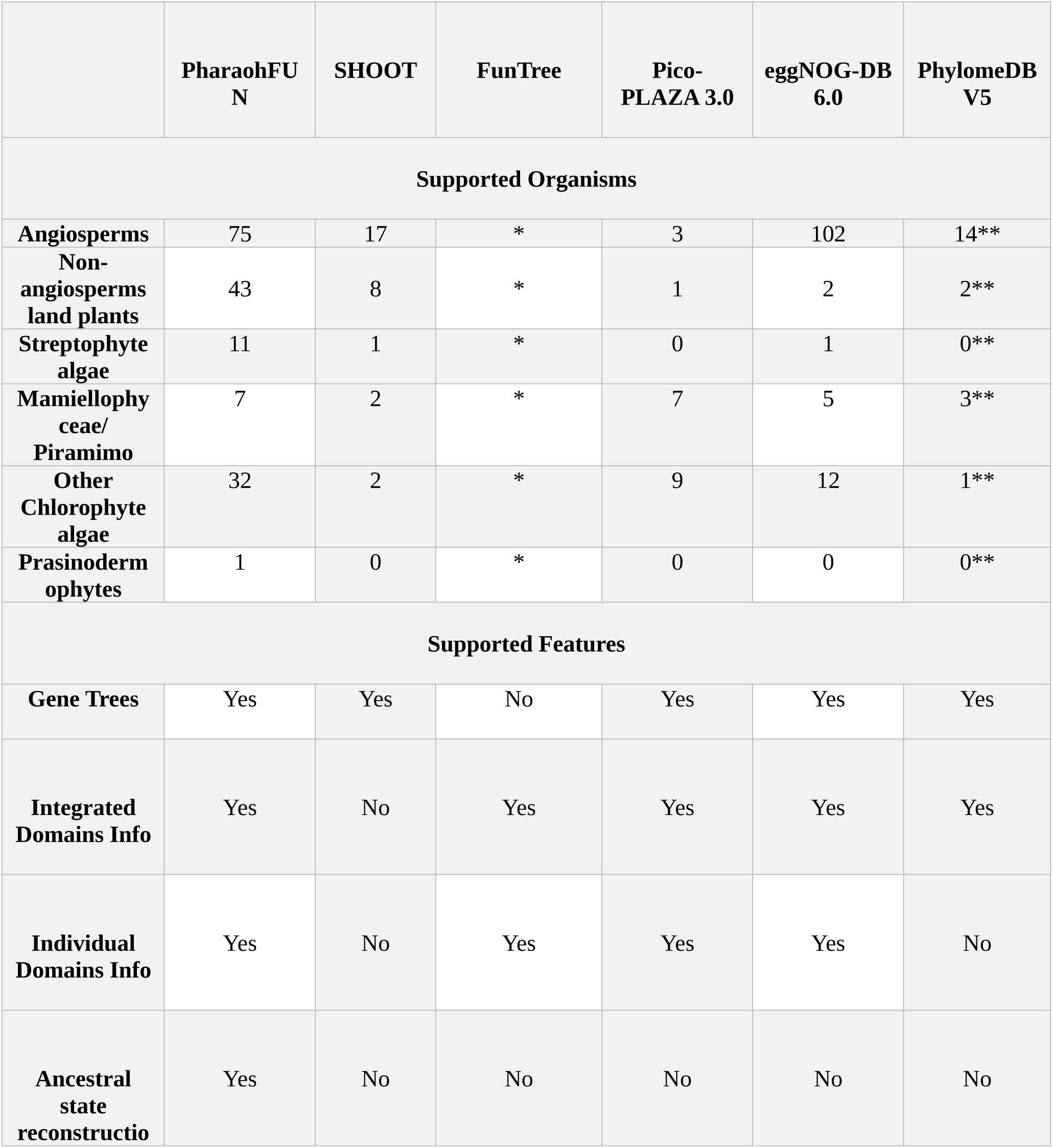
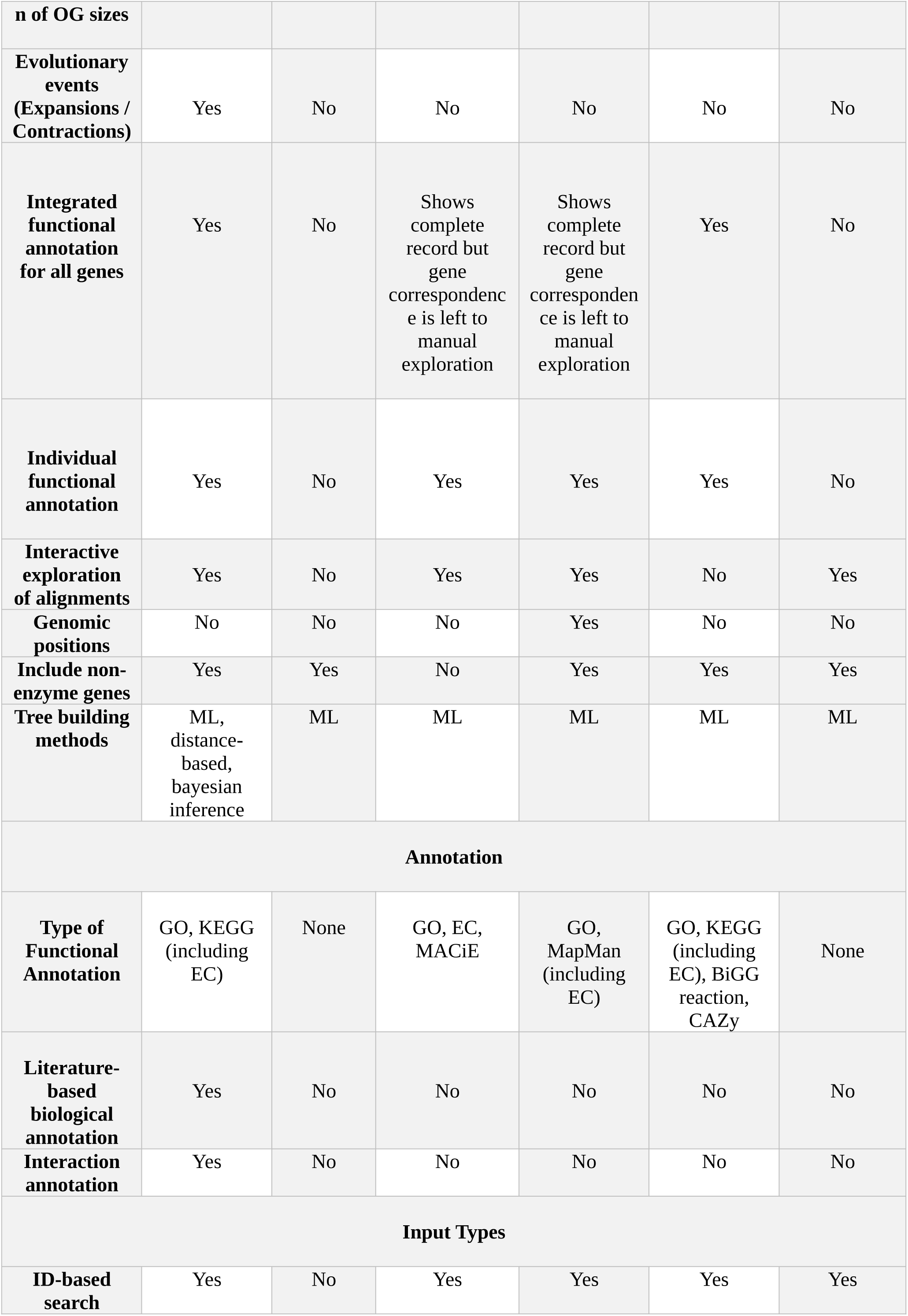
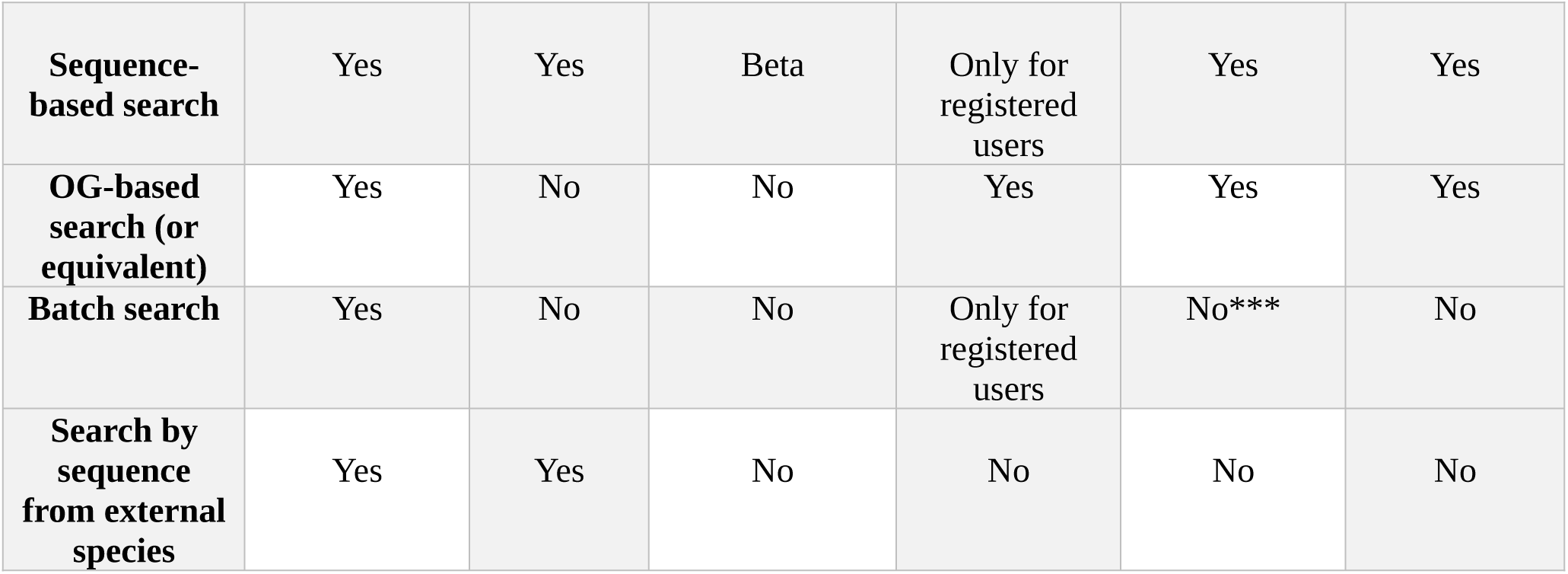
Comparison between phylogenetic web-based tools. *FunTree uses taxonomy data to guide reconciliation of trees, but species tree is not explicitly shown. ** Numbers correspond to the largest plant phylome. *** eggNOG-DB 6.0 does not incorporate a batch mode, but fully automatic functional annotation of sequences can be performed using eggNOG-mapper v2 on eggNOG-DB 5.0.

SHOOT primarily focuses on the phylogenetic location of new genes by incorporating them into a database through MSA. While it provides links to gene information stored in Phytozome, it lacks modules for domain analysis, orthogroup evolution along species trees, and functional annotation of complete orthogroups. The limited representation of algae among its 30 species Plant database (only 4 chlorophytes and 1 streptophyte) is evident when conducting searches, as demonstrated by the example of searching for the CCA1 gene, which only returns results in terrestrial plants at the default tree level (Sup. Fig. S12A) despite its widespread presence across Viridiplantae (A.-M. Linde et al. 2017).

FunTree is designed to study the evolution of enzyme superfamilies by integrating functional, structural and phylogenetic information. Unlike the previous tool, it reconstructs common ancestors of gene groups. Its focus on specific enzyme superfamilies and their activities restricts its applicability and genes that do not code for enzymes in the registered superfamilies are not analyzed. Moreover, reconstructions are performed at the level of the most likely ancestral enzyme activity (in terms of their EC number), not based on OG status. Therefore, features to explore gene evolutionary history are not provided. In the case of the CCA1 example, neither the Uniprot ID search nor the beta sequence-based search function yielded any result.

To some extent, pico-PLAZA pursues a similar goal to PharaohFUN’s but, while PharaohFUN aims to uniformly represent all Viridiplantae groups, pico-PLAZA focuses on chlorophyte and stramenopiles algae. In this sense, although pico-PLAZA targets 11 chlorophyte genera, it lacks representation for streptophyte algae, which would serve as a liaison to the 4 land plants considered in its database, and only supports one non-angiosperm embryophyte species (*P. patens*). The lack of linking elements between these evolutionarily distant groups produces separation into isolated instances. Although this may lead to the creation of refined gene groups, it hinders an integrated analysis of OGs. Bias can be incorporated in the clustering process towards the creation of small highly similar gene groups corresponding to very closely related species excluding related but distant genes from OGs. In contrast, PharaohFUN employs uniform species sampling to eliminate bias towards any specific clade. This approach allows for the implementation of an analysis of the evolution of different OGs along the species tree, a feature not supported by pico-PLAZA. Furthermore, in terms of annotation, PharaohFUN incorporates the new literature annotation utility provided by PlantConnectome, facilitating the exploration of biological data associated with specific terms of interest. It also enables annotation and representation of KEGG pathways, STRING networks, structural models for STRING-supported genes and coevolution of OGs associated with interacting proteins. While SHOOT lacks the functionalities to directly perform functional analysis for a whole OG and pico-PLAZA indirectly allows it (through individual gene search, selection and custom gene set analysis restricted for registered users in Workbench), PharaohFUN overcomes this limitation by providing specific analysis features for these tasks within the same platform, always centered in whole OGs or custom genes and clades. As a comparison with PharaohFUN, the search for the gene encoding CCA1 in *A. thaliana* in pico-PLAZA found an orthologous gene family (ORTHO03P024982) with only four angiosperm clade-specific genes (Sup. Fig. S12B), despite its presence in chlorophyte species such as *O. tauri*, reflecting the aforementioned bias due to lack of evolutionary cohesion (A.-M. Linde et al. 2017). There exist other PLAZA instances focused on specific groups, such as dicots and monocots (Van Bel et al. 2022), but all suffer from the limitation in generalization for evolutionary studies over long distances due to the lack of representatives outside those clades.

This limitation arises from the trade-off between exploring the full evolutionary range of plant evolution and the need to use genomes with sufficient completeness and contiguity for chromosomal position studies (Vandepoele et al. 2013; Van Bel et al. 2022). Thus, while PharaohFUN focuses on the study of proteomes across clades without providing this feature, pico-PLAZA sacrifices this continuity in species sampling in favor of incorporating data from genomic regions, at the cost of not including species that serve as a link between land plants and chlorophyte algae. The impact of this design is accentuated by the fact that land plant species supported by pico-PLAZA have undergone several Whole Genome Duplication events, making it more likely to create OGs more focused on that particular clade, without recovering their algal counterparts (Wang et al. 2022). While consistent with all other instances of PLAZA, this decision effectively leaves unsupported a group of organisms currently heavily represented in evolutionary studies, where omics and molecular studies are already routine (Bierenbroodspot et al. 2024).

eggNOG-DB (Hernández-Plaza et al. 2023) is a popular tool for performing comparative genomic analysis on a wide range of organisms, including Viridiplantae. It is focused on complete OG information, with individualized data available in the tree representation. It includes detailed functional annotation, incorporating GO, KEGG, CAZy and BiGG data. However, it incorporates a very low proportion of non-angiosperm land plants and green algae within its Viridiplantae set, making it difficult to use in plant evolution studies outside of flowering plants. It does not offer a batch mode, but fully automatic large-scale functional annotation of the sequences can be performed via eggNOG-mapper v2 (Cantalapiedra et al. 2021), making it a powerful tool for transferring functional data. However, this analysis does not report gene trees, MSAs or information on the evolutionary history of the sequences. Currently, eggNOG-mapper v2 supports up to version 5.0 of eggNOG-DB. In this case, eggNOG-DB 6.0 did recover the CCA1/LHY ortholog in *O. tauri* (Sup. Fig. S12C).

Finally, PhylomeDB (Fuentes et al. 2022) is another popular tool for phylogenetic studies spanning the tree of life. It contains several collections or phylomes focused on different species and provides information on them such as gene trees, domains and MSAs. However, in general it supports a low number of plant species, as can be seen in Table 2 for its largest plant phylome. PhylomeDB also exhibits the aforementioned limitations in terms of representatives outside the angiosperm group. Thus, despite incorporating *O. tauri* into the collection, it does not recover its CCA1/LHY ortholog, nor any outside the angiosperms clade (Sup. Fig. S12D). Furthermore, this tool is focused on phylogenetic analysis of many species but does not incorporate functional information or evolutionary history reconstruction for them.

With this comparison, it becomes clear that PharaohFUN supports a significantly higher number of plant species than the other tools. The difference is dramatic in the case of non-angiosperm land plants and streptophyte and chlorophyte algae, while maintaining an intensive sampling of angiosperms without compromising the balance of clades, thanks to the creation of a phylodiverse model to which subsequently assign the rest of the genes. PharaohFUN also stands out for its integration of abundant functional and domain information with interactive exploration of MSAs and its flexibility regarding search modes, allowing large-scale evolutionary studies or the inclusion of sequences from species absent in the dataset, features not offered by most tools. Finally, it is the only one that offers information on physical interactions for studies of coevolution between different proteins, literature-based annotation and reconstructions of the evolutionary history of orthogroups along the species tree, in terms of ancestral states at internal nodes and mapping of expansion/contraction/gain/loss events on its branches.

### Conclusions and future directions

In summary, PharaohFUN significantly increases and unifies the evolutionary range for studies outside the flowering plants. It addresses the growing demand for research on photosynthetic eukaryotes outside this dominant group, particularly highlighting its utility in analyzing aquatic species of biotechnological or evolutionary interest and non-seed land plants. In addition, PharaohFUN stands out among other evolutionary analysis tools due to its unique features integrating domain, sequence and functional annotation information into a single platform. Thus, it provides a comprehensive approach that enables researchers to conduct in-depth and holistic analysis of gene evolution and function, making PharaohFUN a valuable tool for the research community on photosynthetic organisms.

As future directions, we expect PharaohFUN to be widely embraced by users in the field of plant sciences due to its user-friendly interface and its ability to facilitate evolutionary and functional analysis across numerous species of interest. The tool will continue to evolve based on feedback and interactions with users being regularly updated to include new representative species from different clades dealing with the rapid expansion in the number of genomes available. PharaohFUN’s modular design allows for seamless incorporation and analysis of new species, without requiring major changes to its underlying architecture. This adaptability ensures that PharaohFUN will remain a relevant and effective tool meeting the evolving needs of the scientific community working in photosynthetic organisms.

## Acknowledgments

We would like to acknowledge the revision and testing of the tool carried out by several members of the “Spanish National Research Network on Evolution of Regulatory Mechanisms in Development and Plant Signalling” which include but are not limited to Pilar Cubas, Miguel Ángel Blázquez, Javier Fuentes and Lluís García.

## Author Contributions

FJRC and MGG conceived and conceptualized the project. FJRC, MGG, MRG and JHG designed the structure of the tool. MRG, VRG, ESP and FJRC developed the web app. MRG, VRG, ESP, CA, FJRC and JHG carried out the collection of genomes and annotation of photosynthetic eukaryotes included in the tool. MRG, FJRC and MGG performed all the analysis and interpreted the results. MRG, FJRC and MGG wrote the manuscript. All authors have read and approved the manuscript.

## Data Availability

The complete source code for PharaohFUN is freely available at the following GitHub repository: https://github.com/ramosgonzmarc/PharaohFUN

## Figures

**Sup. Figure S1.** PharaohFUN’s complete species tree. Dated species tree used for model construction. **Left**: dated species tree measured in millions of years. The inferred gene duplication number of internal nodes is mapped by circle color and size. **Right**: gene duplication number of extant species. Clades are distinguished by different bar color according to legend.

**Sup. Figure S2.** Gene tree corresponding to the complete CCA1 OG. Three differentiated clades are highlighted.

**Sup. Figure S3.** PFAM domains location in sequences from the complete CCA1 OG.

**Sup. Figure S4.** Results for the WRKY family. **A.** WRKY OG tree. Notation of groups I and II followed that of (Huang et al. 2012) and notation of group III that of (Wang et al. 2015). **B.** Evolutionary events in Viridiplantae for the OG comprising TIR-NB-LRR proteins.

**Sup. Figure S5.** OGs containing C2H2 genes in *A. thaliana* mapped onto the AtZFPs tree adapted from (Li et al. 2022).

**Sup. Figure S6.** Heatmap showing the row-normalized gene number per OG associated to TF families in 54 Viridiplantae species.

**Sup. Figure S7.** Species tree with the number of OGs associated to TF families in *A. thaliana* exhibiting its MRCA in each of its internal nodes. Color and point size determine total number.

**Sup. Figure S8.** Species tree with the number of gained OGs associated to TF families in *A. thaliana* in each of its internal and terminal nodes. Color and point size determine total number.

**Sup. Figure S9.** Species tree with the number of expanded OGs associated to TF families in *A. thaliana* in each of its internal and terminal nodes. Color and point size determine total number.

**Sup. Figure S10.** Species tree with the number of contracted OGs associated to TF families in *A. thaliana* in each of its internal and terminal nodes. Color and point size determine total number.

**Sup. Figure S11.** Species tree with the number of lossed OGs associated to TF families in *A. thaliana* in each of its internal and terminal nodes. Color and point size determine total number.

**Sup. Figure S12.** Results for CCA1 rendered by **A.** SHOOT, **B.** pico-PLAZA 3.0, **C.** eggNOG-DB 6.0 and **D.** PhylomeDB.

## Tables

**Sup. Table S1.** Genomic data used in PharaohFUN, including species names, genome version and genome source.

**Sup. Table S2.** Constraints used for dating the species tree in Mya. Restrictions are indicated by minimum and maximum time points for different ancestors of the clades, using fossil data from (Strassert et al. 2021) and data from (Kumar et al. 2022).

**Sup. Table S3.** ExaBayes parameter file.

**Sup. Table S4.** Number of genes associated to each OG of those interacting with CCA1 orthologs in *A. thaliana*, *P. patens* and *O. tauri*.

**Sup. Table S5.** Assignment of OGs by TF family, their common ancestor and the number of genes in each species according to PharaohFUN results.

**Sup. Table S6.** Number and type of PFAM domains of *A. thaliana* genes in each TF-related OG according to PharaohFUN results.

**Sup. Table S7.** Evolutionary events occurred in each node from the species tree for TF-related OGs.

**Sup. Table S8.** Sensitivities per TF family in each test species using *A. thaliana* genes as source.

**Sup. Table S9.** Sensitivities per TF family in each test species using *A. thaliana*, *B. prasinos*, *V. carteri*, *P. patens* and *S. bicolor* genes as source.

