## Supplementary material for "PharaohFUN: PHylogenomic Analysis foR plAnt prOtein History and FUNction elucidation": Supp: Sup_TableS3.txt

#NEXUS

[ In this config file everything is commented out.  Remove the square
  brances to activate an option. ]

[ In most instances, it is convenient to use the scientific notations also
   for integer variables (e.g., write 1e6 for requesting numgen=1000000) ] 

[ all keywords are case-insensitive ]


[================================================================]
[                               GENERAL                          ] 
[================================================================]
begin RUN; 
 [ RUN PARAMETERS ]
  numRuns  2  	   	[ number of independant runs ]
  numGen 50000  		[ number of generations to run. If a convergence criterion is used, this is the minimum number of generations to run.  ] 
 [ diagFreq 5000 ] 		[ check for convergence of multiple independent runs after this many generations ]
 [ samplingFreq 500 ] 	        [ take a sample every n generations ]
 [ tuneFreq 100 ]    	 	[ tune move parameters every n generations, set to 0, if you want to disable parameter tuning ] 
 [ printFreq 1000 ]  		[ print a sample to the screen every n generations, set to 0 if you want to disable output info about the state of the chains  ]

  parsimonyStart true    [ if true: use parsimony trees as starting trees, else: use random starting trees ] 
 [ heatedChainsUseSame  false ]   [ should heated chains start with the same tree as the cold chain? default: false] 
 [ checkPointInterval   10000 ]	  [ defines how often a checkpoint is written ]

 [ MCMCMC ]
  numCoupledChains  2 	[ number of chains per independent run (only one is cold)  => must be > 0] 
 [ heatFactor 0.1 ]     	[ the heat increment added to each hotter chain (this is not the inverse heat)   ]
 [ numSwapsPerGen 1 ]           [ number of swaps attempted on average per generation ] 

 [ CONVERGENCE ]
 [ convergenceCriterion mean ]  [ indicates whether a topological convergence criterion should be used. Valid values: none, mean (default), max ] 
 [ sdsfIgnoreFreq  0.1 ]        [ ignore clades for which the relative frequency does not exceed this value in any chain (here 10%) ]
  sdsfConvergence 0.05        [ consider runs converged (and stop) once the asdsf or msdsf is below this value (here 1%): 1-5% is considered good, 0.5% can be considered very good convergence ] 

 [ BURNIN ] 
 [ burninGen 2000 ]		    [ exact number of generations that are discarded for diagnostics ]  
 [ burninProportion      0.25 ]     [ discard this proportion of all initial samples as burnin  ]

 [ proposalSets     false ]      [ enables accumulation of proposal into proposalSets. Very IMPORTANT, if you run an analysis using multiple unlinked parameters ]
end; 


begin params; 
[================================================================]
[ PARAMETERS: definition and linking ]
[================================================================]

 [ stateFreq = (0) ]       [ state frequencies ] 
 [ rateHet   = (0) ]       [ rate heterogeneity among sites ] 
 [ revMat    = (0) ]       [ reversible substitution rate matrix ] 
 [ brlens    = (0) ]       [ branch lengths ] 
 [ aaModel   = (0) ]      [ fixed-rate amino acid substitution matrix ] 

 brlens    = (0)       [ branch lengths are now linked by default. Use this statement to UNLINK branch lengths (assuming, you have 4 partitions) ] 

[ syntax for linking: ] 
[ brlens = (0-3)      ]    [ linked: one branch length parameter, linked across all 4 partitions ]
[ brlens = (0:3)      ]    [ unlinked: 4 branch length parameters (one for each partition)  ] 
[ brlens = (0+1,2,3)  ]    [ the first parameter is linked for partitions 0 and 1 ] 
[ brlens = (0-2,3)    ]    [ first parameter linked across the first 3 partiions, a separate parameter for the last partition ] 
  
end;  


[================================================================]
[ PRIOR configuration ]
[================================================================]
begin PRIORS; 
[ TOPOLOGY: fixed or uniform  ]
[   topoPr fixed()      ] 
[   topoPr uniform(0,0) ]


[ BRANCH LENGTHS: uniform, exponential or fixed ]
[ brlenpr uniform(1e-6,4)     ] 
[ brlenpr fixed()             ]      [ you should have provided a user tree with branch lengths, otherwise 0.1 ]  
[ brlenpr fixed(0.1)          ]      [ fixes branch lengths to 0.1 ] 
[ brlenpr exponential(10)     ]      [ exponential distribution with mean value 0.1 ] 

[ SUBSTITUTION MATRIX: dirichlet or fixed, 6 values for DNA, 190 for AA (you must have defined a revMat parameter for the respective partitions) ]
[ order of the substitution rates corresponds to the order in the ExaBayes_parameters* output file ]
[ revMatPr dirichlet(1,2,3,
		       4,5,
		         6) ]            [ for DNA (rates are relative to each other and will be normalized, s.t. they sum up to 1) ]
[ revMatPr dirichlet( 1.0,1.0,1.0,1.0,1.0,1.0,1.0,1.0,1.0,1.0,1.0,1.0,1.0,1.0,1.0,1.0,1.0,1.0,1.0,
  	   	          1.0,1.0,1.0,1.0,1.0,1.0,1.0,1.0,1.0,1.0,1.0,1.0,1.0,1.0,1.0,1.0,1.0,1.0,
			      1.0,1.0,1.0,1.0,1.0,1.0,1.0,1.0,1.0,1.0,1.0,1.0,1.0,1.0,1.0,1.0,1.0,
			          1.0,1.0,1.0,1.0,1.0,1.0,1.0,1.0,1.0,1.0,1.0,1.0,1.0,1.0,1.0,1.0,
				      1.0,1.0,1.0,1.0,1.0,1.0,1.0,1.0,1.0,1.0,1.0,1.0,1.0,1.0,1.0,
				          1.0,1.0,1.0,1.0,1.0,1.0,1.0,1.0,1.0,1.0,1.0,1.0,1.0,1.0,
					      1.0,1.0,1.0,1.0,1.0,1.0,1.0,1.0,1.0,1.0,1.0,1.0,1.0,
					          1.0,1.0,1.0,1.0,1.0,1.0,1.0,1.0,1.0,1.0,1.0,1.0,
						      1.0,1.0,1.0,1.0,1.0,1.0,1.0,1.0,1.0,1.0,1.0,
						          1.0,1.0,1.0,1.0,1.0,1.0,1.0,1.0,1.0,1.0,	
							      1.0,1.0,1.0,1.0,1.0,1.0,1.0,1.0,1.0,
							          1.0,1.0,1.0,1.0,1.0,1.0,1.0,1.0,
								      1.0,1.0,1.0,1.0,1.0,1.0,1.0,
								          1.0,1.0,1.0,1.0,1.0,1.0,
									      1.0,1.0,1.0,1.0,1.0,
									          1.0,1.0,1.0,1.0,
										      1.0,1.0,1.0,
										          1.0,1.0,
											      1.0 ) ]     [ 189 values for AA]
[ revMatPr fixed(1,2,3,4,5,6) ]                [ for DNA, rates will be normalized  ] 

[ STATE FREQUENCIES: dirichlet or fixed, 4 values for DNA, 190 for AA (you must have defined a stateFreq parameter for the respective partitions) ] 
[ order of the frequecies corresponds to the order in the ExaBayes_parameters* output file ]
[  statefreqpr  dirichlet(0.25,0.25,0.25,0.25)  ]                                                           [ for DNA,  could also be relative rates again  ]
[  statefreqpr dirichlet(1.0,1.0,1.0,1.0,1.0,1.0,1.0,1.0,1.0,1.0,1.0,1.0,1.0,1.0,1.0,1.0,1.0,1.0,1.0,1.0) ] [ for AA  ] 
[  stateFreqPr fixed(1,2,3,4)                   ] 

[ RATE HETEROGENEITY: alpha parameter of gamma distribution, exponential, fixed or uniform ]
[ shapePr exponential(2) ] 
[ shapePr uniform(0.0000001,200) ] 
[ shapePr fixed(1)   ] 

[ fixed-rate MODELS for AMINO ACID partitions, fixed or specified ] 
[ aaPr disc( WAG=3, REMAINDER=1 ) ]            [ high prior belief in WAG, all remaining models have a lower prior belief ]
[ below a complete list of AA models implemented in ExaBayes ] 
[ aaPr disc(  DAYHOFF=1, DCMUT=1, JTT=1, MTREV=1, WAG=1, 
       	      RTREV=1, CPREV=1, VT=1, BLOSUM62=1, MTMAM=1, 
	      LG=1, MTART=1, MTZOA=1, PMB=1, HIVB=1, HIVW=1, 
	      JTTDCMUT=1, FLU=1) ]   
[ aaPr fixed(WAG) ]                            [ fix the model to WAG ] 

end; 


 [================================================================]
 [ PROPOSAL configuration ]
 [================================================================]
begin proposals; 
 [ each proposal has a relative weight, use 0 to deactivate ] 

 [ for disabling a category, please set it to fixed in the prior
   section rather than to disable all proposals ]
 
  [ TOPOLOGY proposals  ]
  [ eTBR         0 ]       
  [ eSPR         6 ] 
  [ stNNI        6 ] 
  [ parsimonySPR 6 ] 
  [ likeSpr      2 ]		[ new posterior guided SPR ]

  [ BRANCH LENGTHS proposals ] 
  [ branchMulti    7 ]
  [ treeLengthMult 1  ] 
  [ nodeSlider     0  ] 
  [ blDistGamma    7  ]         [ efficient Newton-Raphson-based branch length proposal ]
  [ biasBLMult     0  ]         [ branch length multiplier that preferably samples short branch lengnths ]

  [ RATE HETEROGENEITY ]
  [ rateHetMulti   1  ] 

  [ SUBSTITUTION RATES  ] 
  [ revMatSlider       0.5 ]    [ for DNA ]
  [ revMatDirichlet    0.5 ]    [ for DNA ] 
  [ revMatRateDirich    2  ]    [ for AA  ]

  [ [ STATE FREQUENIES ]	]	
  [ frequencySlider    0.5  ] 
  [ frequencyDirichlet 0.5  ] 

  [ fixed-rate AA matrices ]    
  [ aaModelJump        1 ] 


  [ PARAMETERS for various moves ]
  [ eSprStopProb 0.5   ]            [ stopping probability for eSPR moves ] 
  [ etbrStopProb 0.5   ]            [ stopping probability for eTBR moves ] 
  [ parsimonyWarp 0.10 ]            [ warp factor for parsimony based moves ]  
  [ parsSPRRadius  10  ]            [ rearrangement radius (around current position) for parsimony SPR ] 

  [ likesprmaxradius 5  ]           [ a radius for the posterior-guided SPR (likeSPR) move: when pruning a subtree, the likeSpr will check all possible re-attachment positions within this radius (expensive!)  ]
  [ likesprwarp 1.0     ]           [ warp factor for posterior-guided SPR move ]
  [ moveOptMode 0       ]           [ values in (0,4), proposes branches simultaneously with any of the topological moves  ] 
  [ useMultiplier true	]           [ in case moveOptMode > 0, use a branch length multiplier instead of a gamma proposal ]

  [================================================================]

end ; 
