## Supplementary figures and images for "PharaohFUN: PHylogenomic Analysis foR plAnt prOtein History and FUNction elucidation"

### Sup_FigS1.png

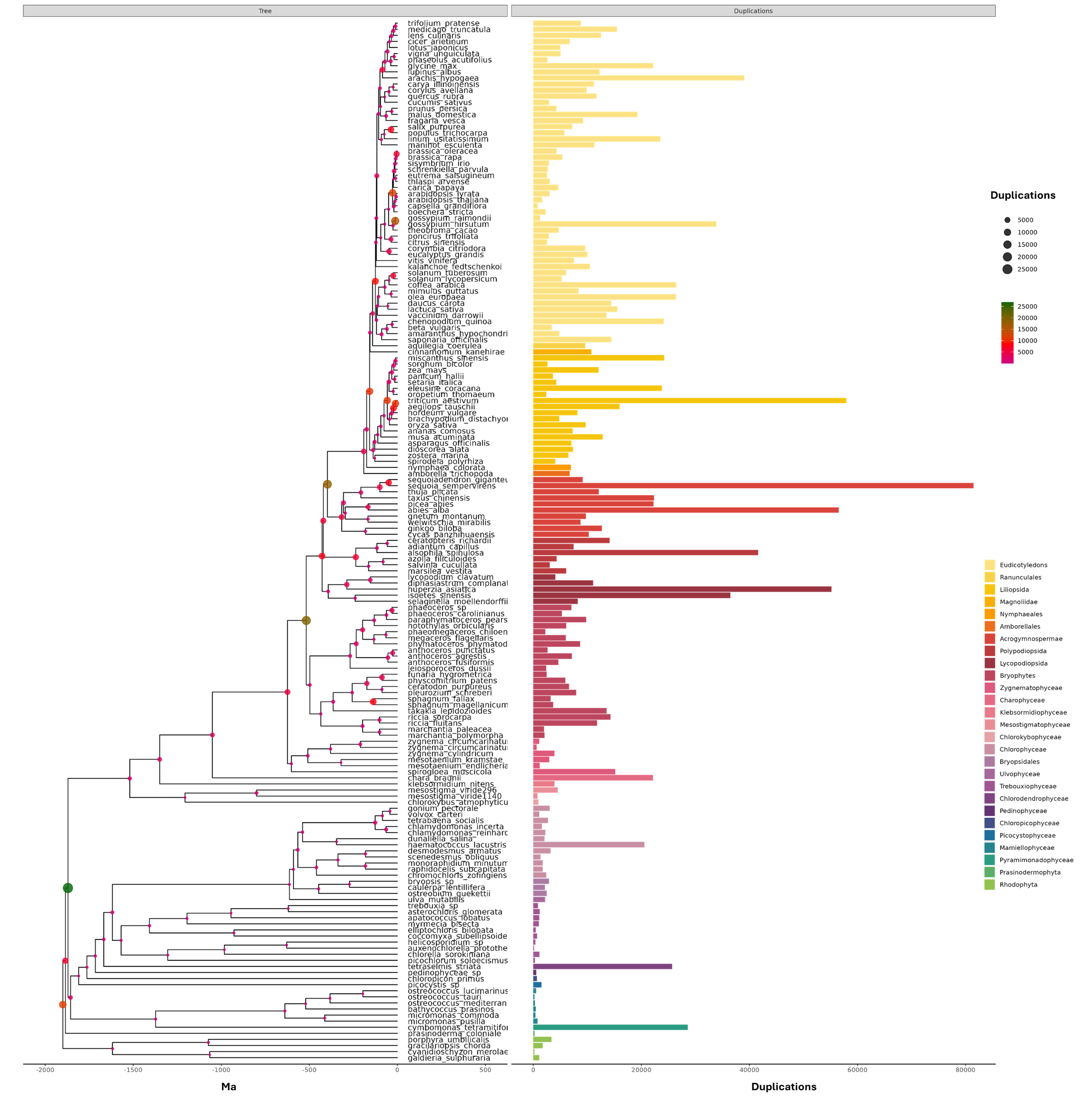

### Sup_FigS2.png

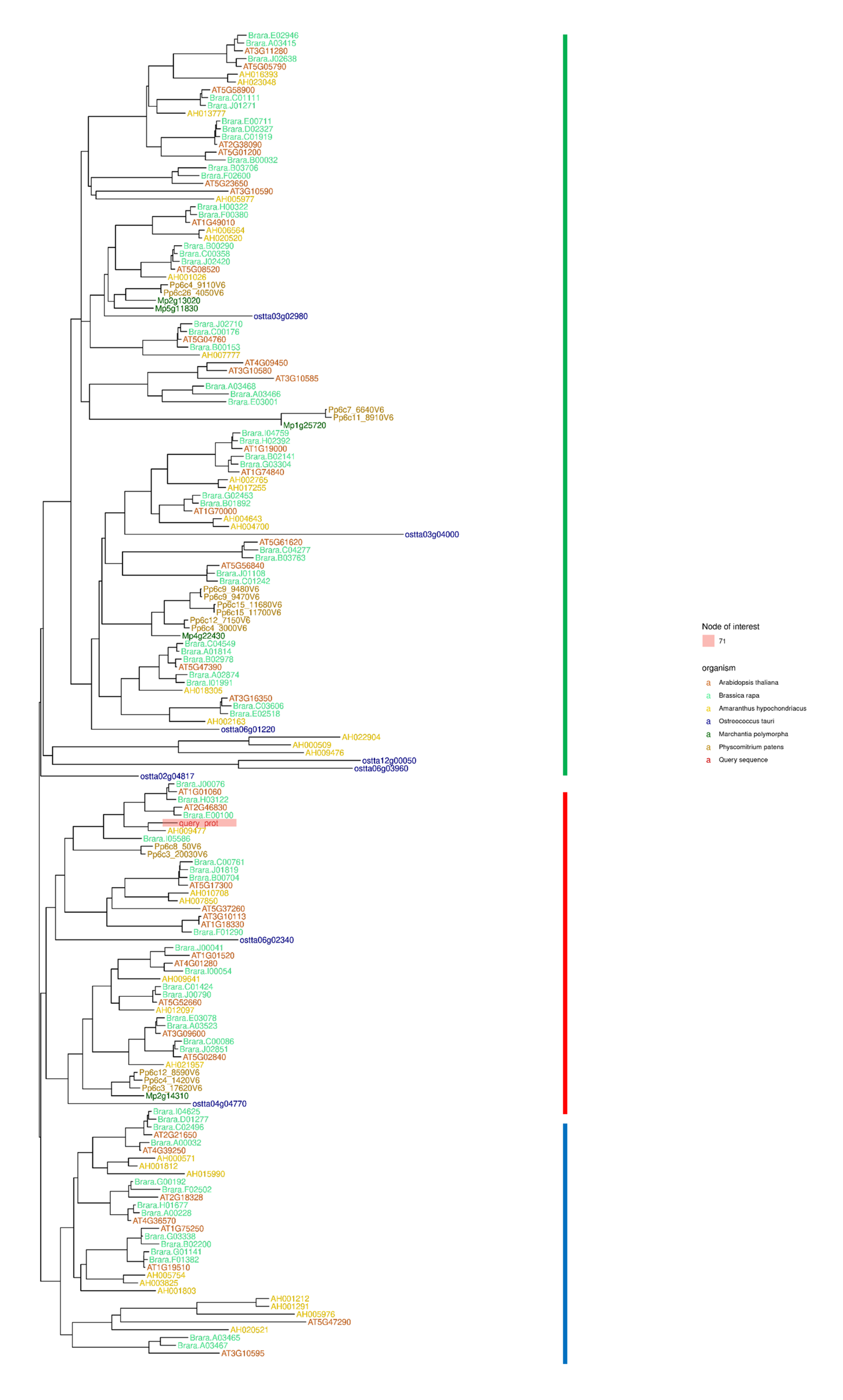

### Sup_FigS3.png

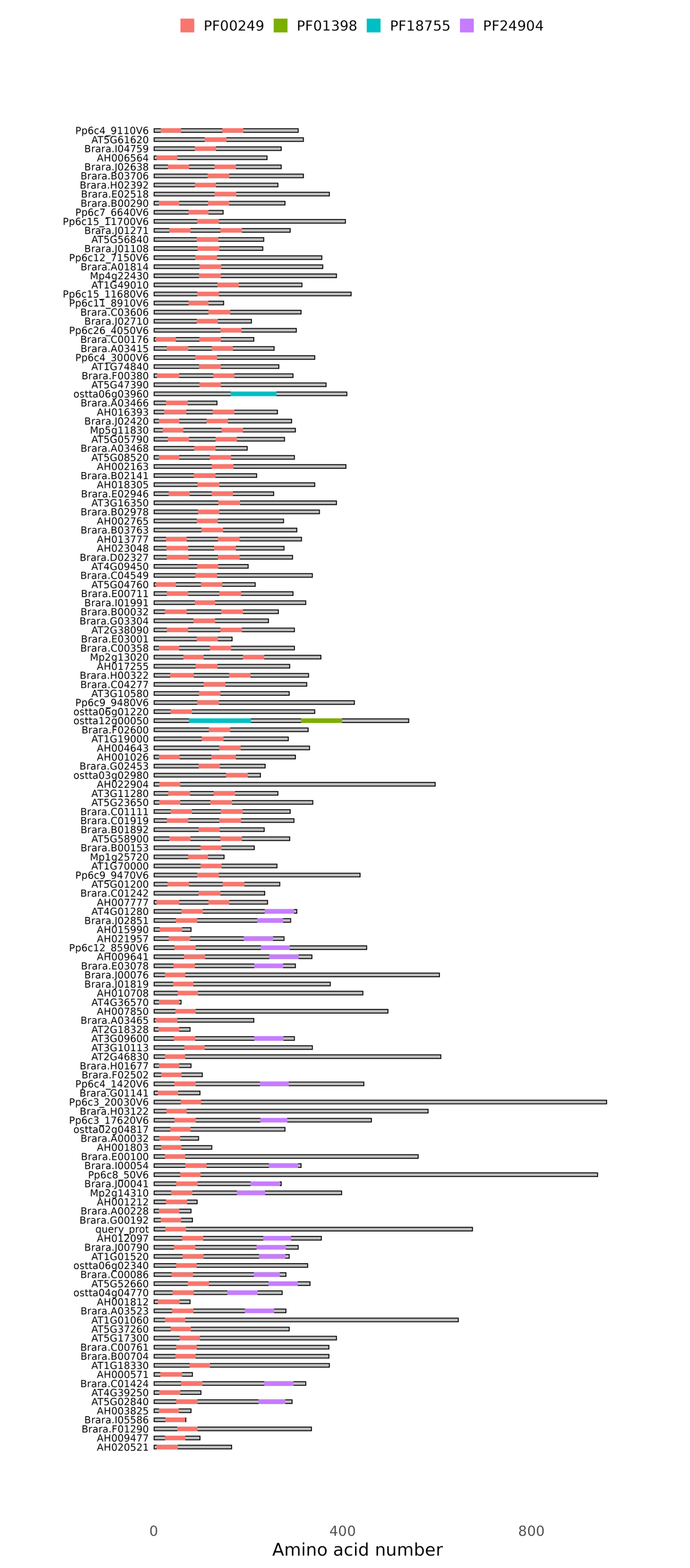

### Sup_FigS4.png

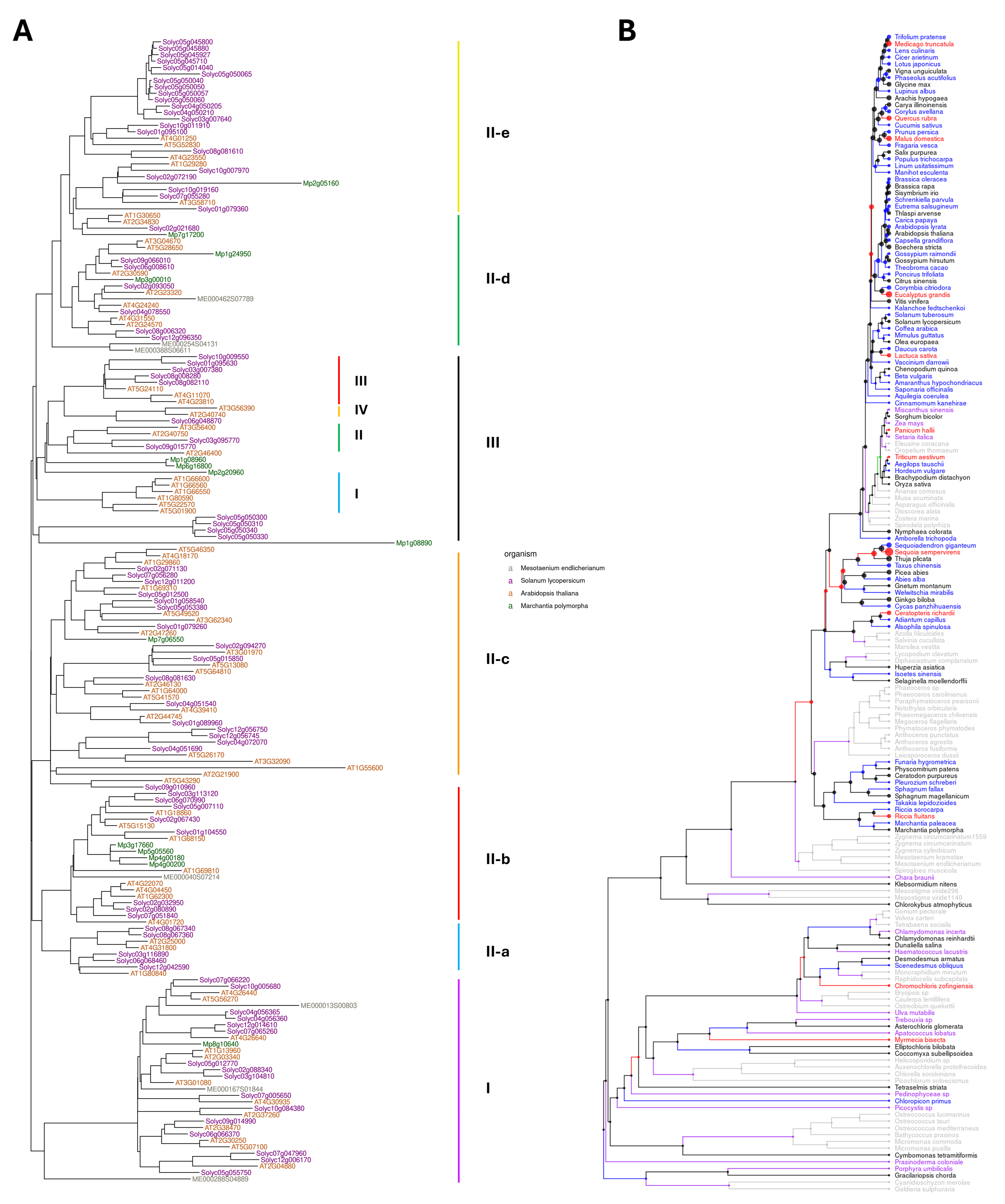

### Sup_FigS5.png

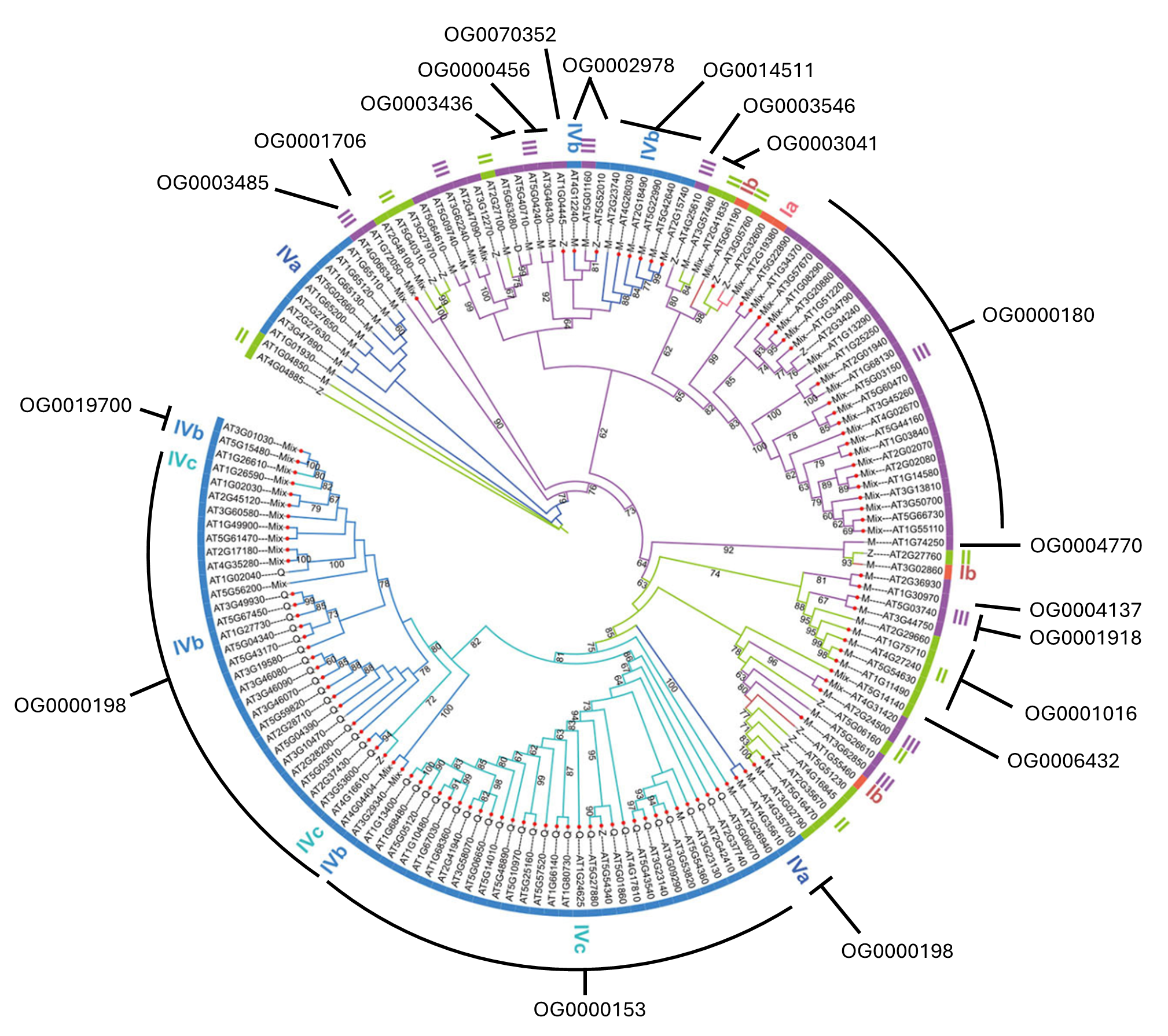

### Sup_FigS6.png

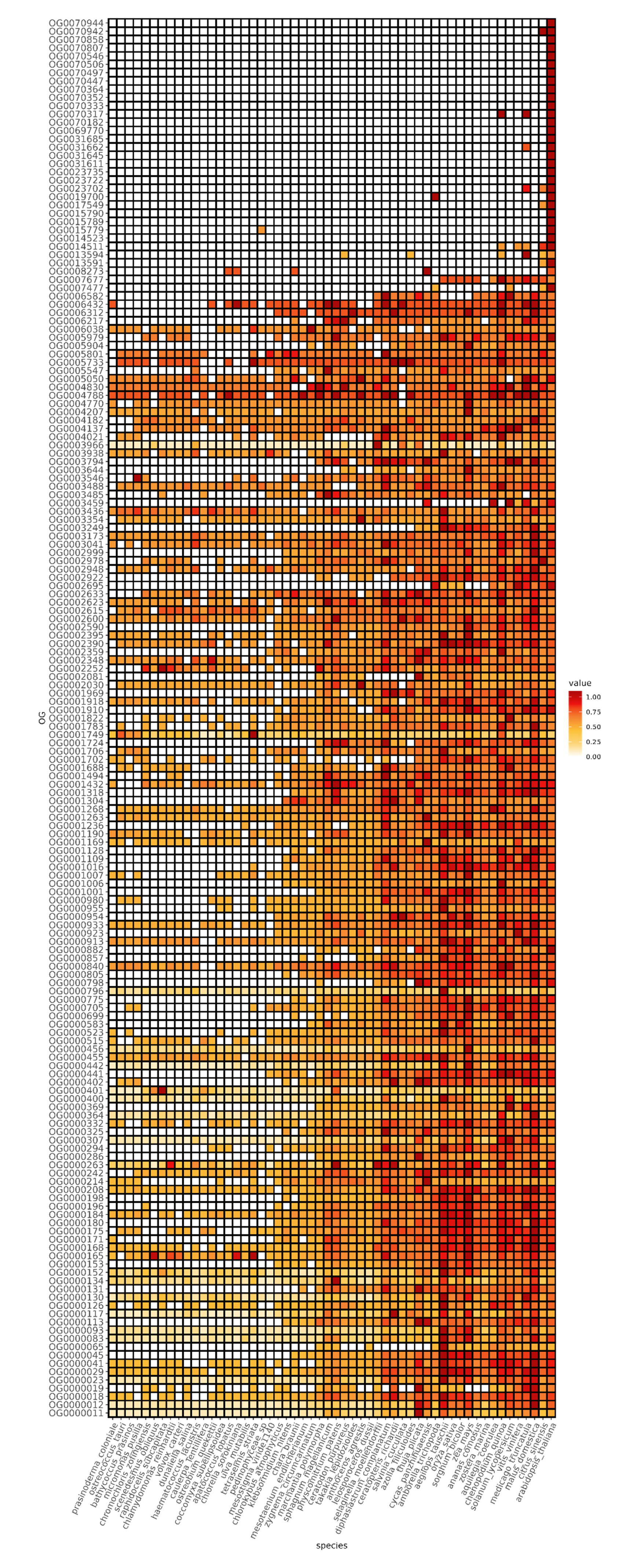

### Sup_FigS7.png

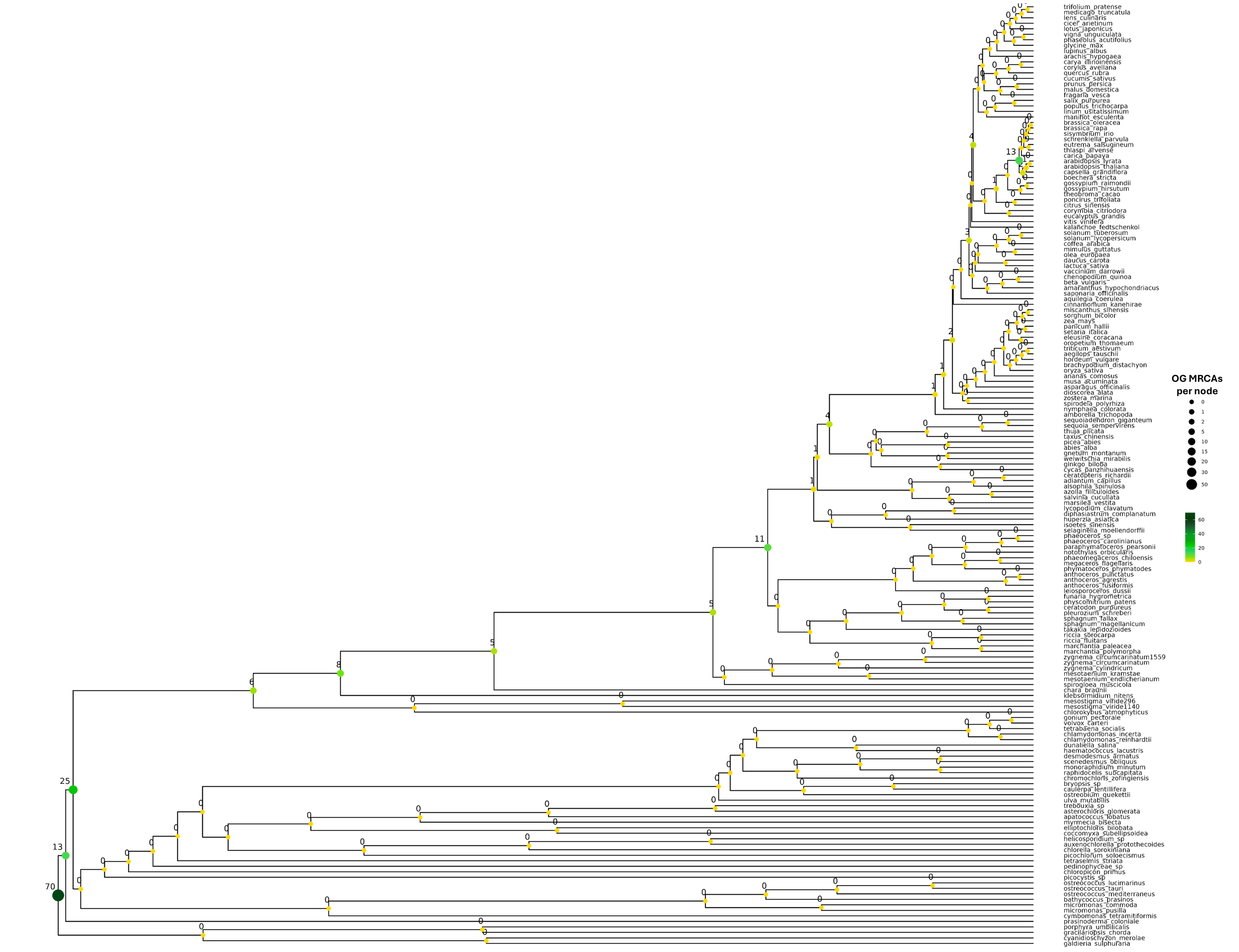

### Sup_FigS8.png

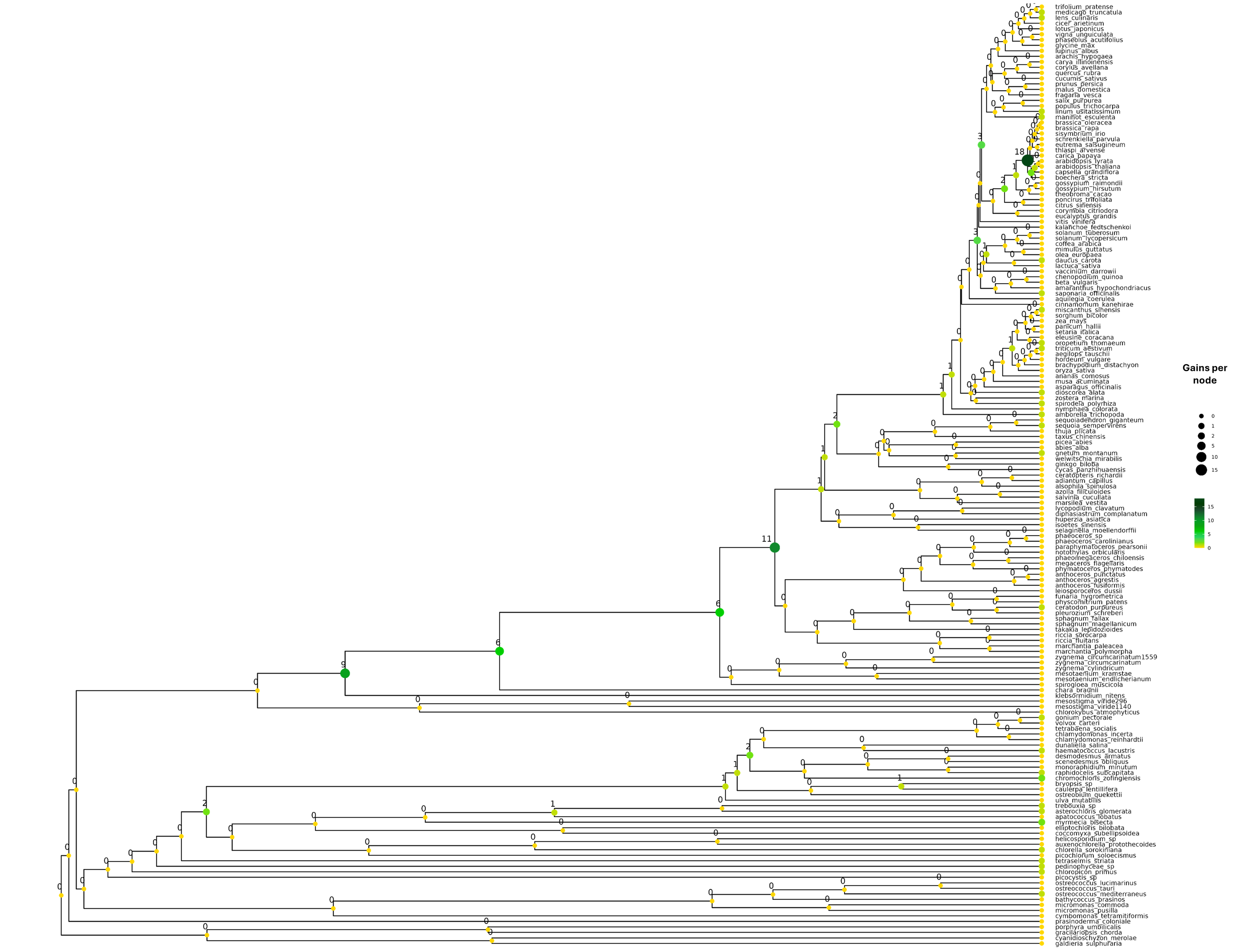

### Sup_FigS9.png

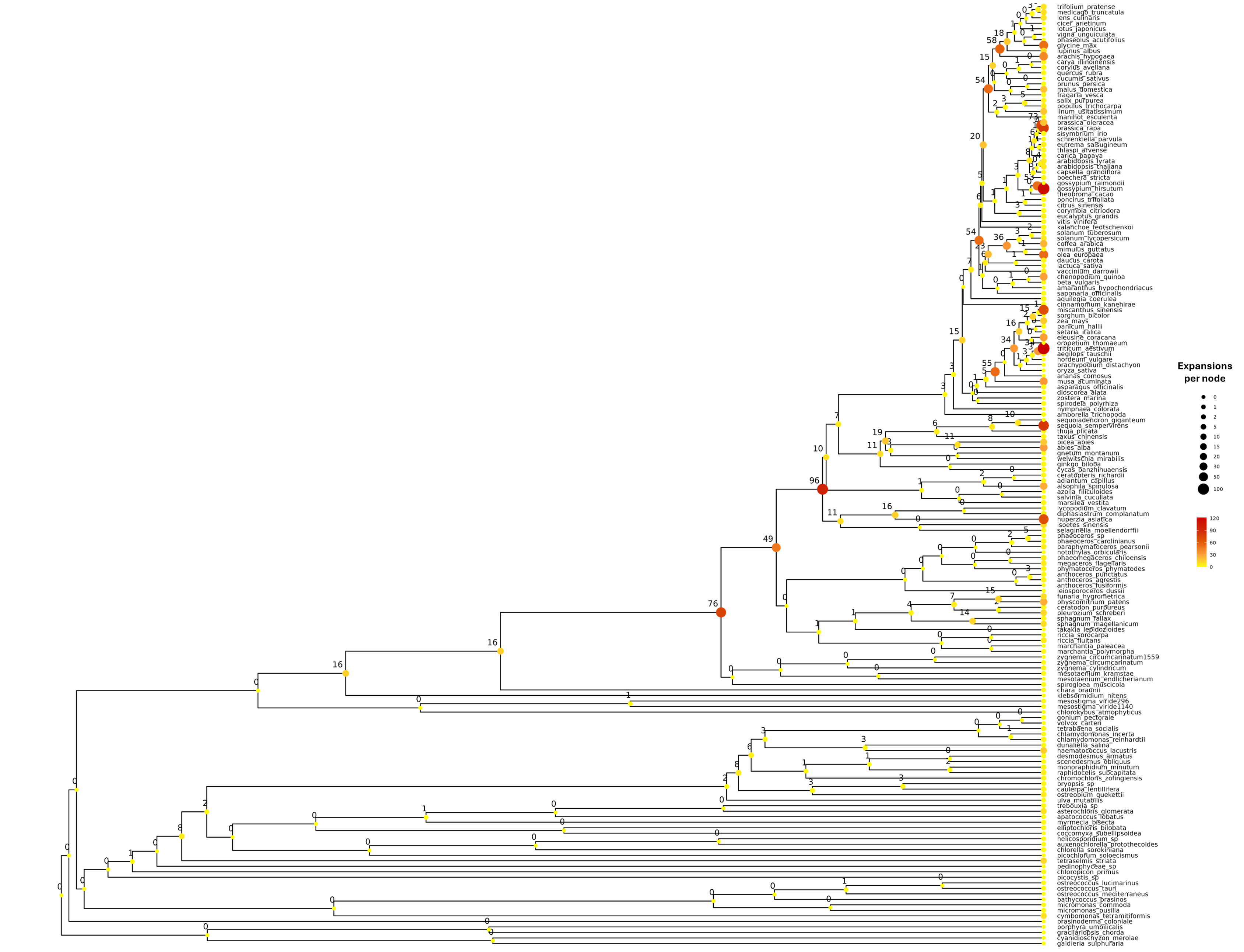

### Sup_FigS10.png

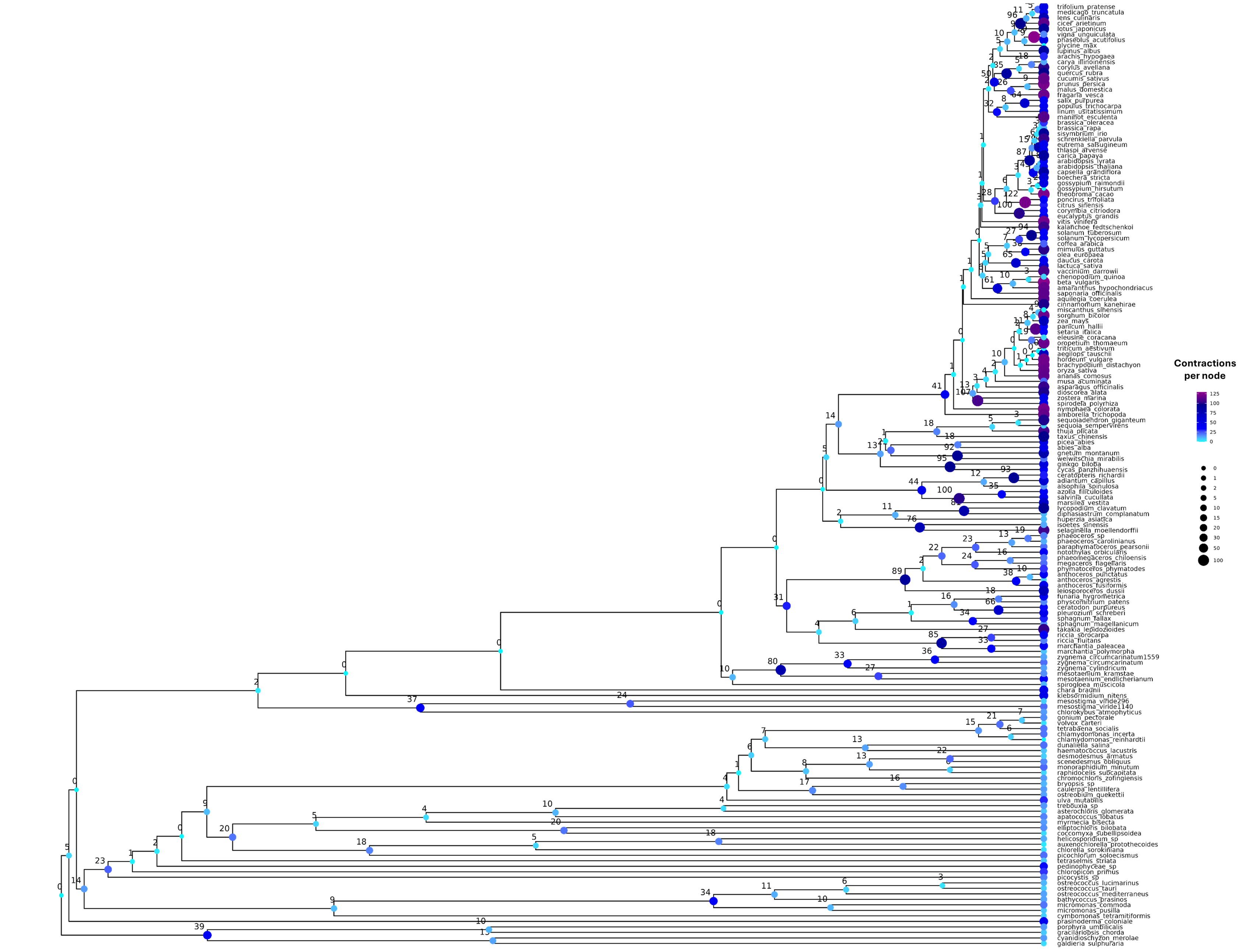

### Sup_FigS11.png

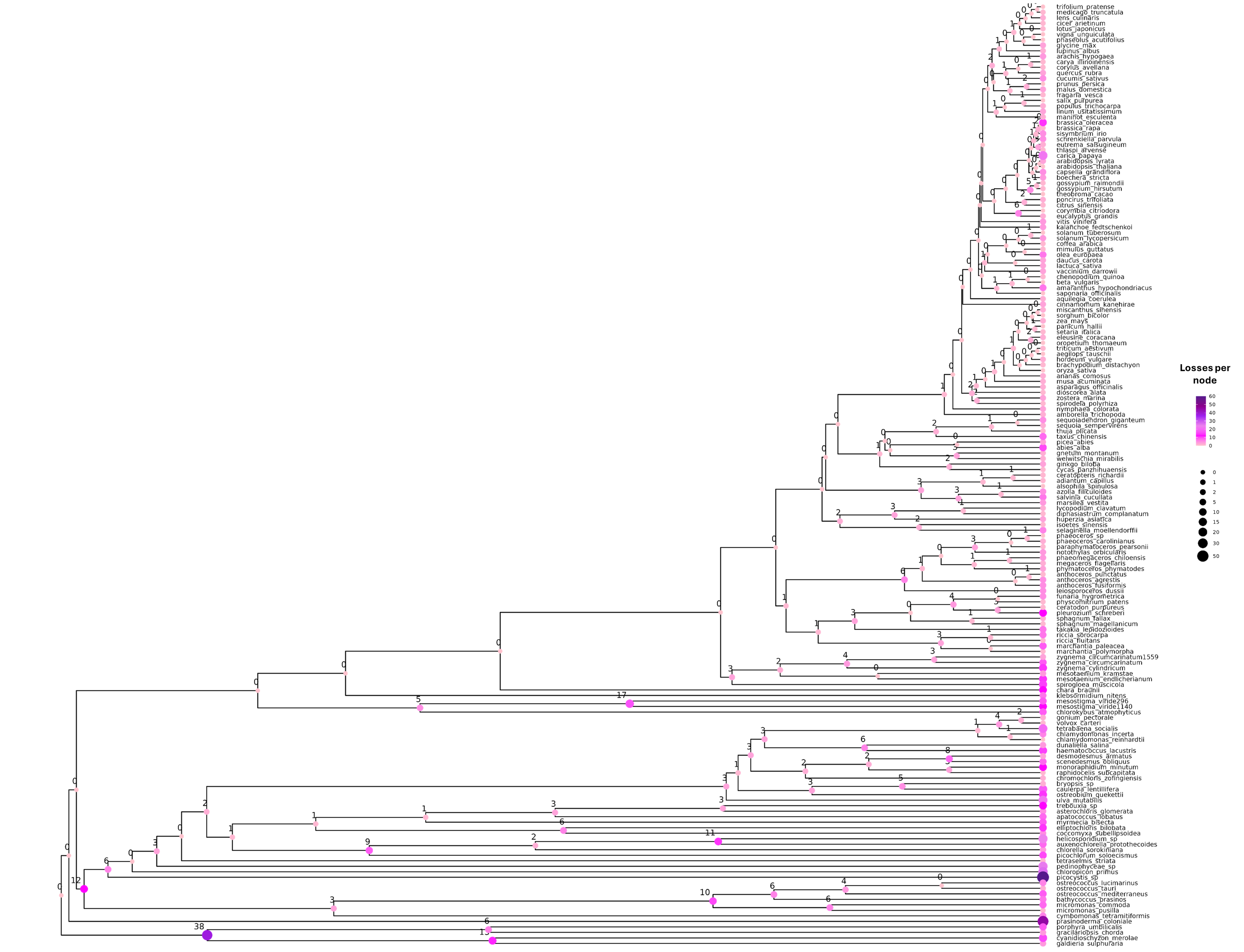

### Sup_FigS12.png

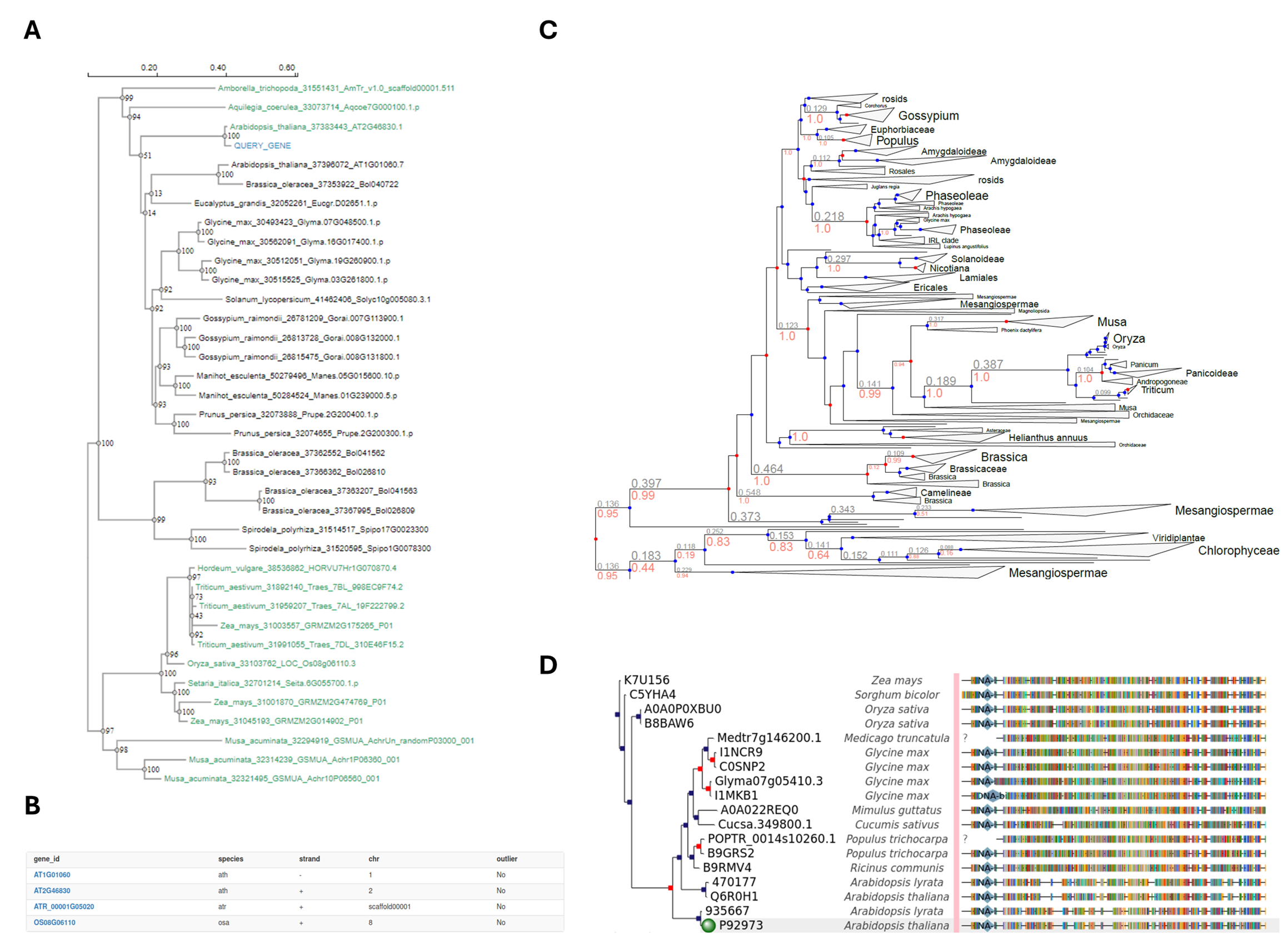
